# Human infants’ understanding of social imitation: Inferences of affiliation from third party observations

**DOI:** 10.1101/050385

**Authors:** Lindsey J. Powell, Elizabeth S. Spelke

## Abstract

Imitation is ubiquitous in positive social interactions. For adult and child observers, it also supports inferences about the participants in such interactions and their social relationships, but the origins of these inferences are obscure. Do infants attach social significance to this form of interaction? Here we test 4- to 5.5-month-old infants’ interpretation of imitation, asking if the imitative interactions they observe support inferences of social affiliation across 10 experimental conditions that varied the modality of the imitation (movement vs. sound), the roles of specific characters (imitators vs. targets), the number of characters in the displays (3 vs. 5), and the number of parties initiating affiliative test events (1 vs. 2). These experiments, together with one experiment conducted with 12-month-old infants, yielded three main findings. First, infants expect that characters who engaged in imitation will approach and affiliate with the characters whom they imitated. Second, infants show no evidence of expecting that characters who were targets of imitation will approach and affiliate with their imitators. Third, analyzing imitative interactions is difficult for young infants, whose expectations vary in strength depending on the number of characters to be tracked and the number of affiliative actors to be compared. These findings have implications for our understanding of social imitation, and they provide methods for advancing understanding of other aspects of early social cognitive development.

## Introduction

Human infants face the basic but critical task of learning about the social beings around them. Who is related to whom, and how? The task of learning about social beings may be especially difficult, because much social behavior—both gestures and speech— depends in part on arbitrary, conventional relations that must be learned (e.g., the words and hand gestures that signal the start of an interaction (“hi!”; a waving hand) and its end (“bye bye!” a flapping hand). Some social behavior, however, relies not on the specifics of the actions but on the relationship between the social partners’ behavior. One such behavior is the imitation, by one party, of another person’s action. Imitation is a universal language for expressing social engagement, because one can only systematically imitate the behavior of another person if one is attending to that person. If the imitated behavior serves no clear instrumental function, moreover, then its performance suggests that the imitator is motivated not only to attend to the target of imitation but to align with the target for social or communicative purposes. Here we use studies of young human infants to probe the origins and nature of the social and communicative functions of imitation, asking how infants interpret imitative interactions they observe as third parties. Do young infants use patterns of imitative behavior to attribute social motives to the partners in that interaction? In particular, do infants attribute prosocial motives to imitators, expecting imitators to affiliate with the targets of their imitation?

### Prosocial imitation and development

Adults often signal their attention to and understanding of another person’s speech by imitating the person’s words and intonation either directly (“OK, two espressos”) or indirectly (“Copy that.”) Even in the absence of such overt communicative motives, however, imitation and mimicry are common, spontaneous components of social interaction that both reflect and elicit liking and prosocial behavior in adults (Bernieri, 1988; Chartrand & Bargh, 1999; Lakin & Chartrand, 2003; Sinclair, Lowery, Hardin & Colangelo, 2005) and children (Thelen, Dollinger & Roberts, 1975; Kinzler, Corriveau & Harris, 2011). Children and adults appear to use imitation and mimicry as social tools, as they increase their copying behavior in the presence of desirable social partners or when the threat of ostracism enhances the drive to affiliate (Lakin, Chartrand & Arkin, 2008; Williams, Cheung & Choi, 2000; Over & Carpenter, 2009; Watson-Jones, Whitehouse & Legare, 2016). In third party contexts, children and adults who witness acts of imitation make a variety of inferences about others’ characteristics and relationships. For example, adults find those who mimic friendly rather than condescending social partners to be more competent, and those who mimic honest rather than dishonest social partners to be more trustworthy (Kavanagh, Suhler, Churchland & Winkielman, 2011; Kavanagh, Bakhtiari, Suhler, Churchland, Holland et al., 2013). Five-year-old children infer social attitudes from imitation, judging an imitator to like the individual she copied more than an individual she chose not to copy (Over & Carpenter, 2015). By middle childhood, therefore, participation in imitation is accompanied by an intuitive, likely implicit, conception of imitation as a social gesture. To date, however, the origins of this conception are unknown.

A tendency to imitate and to respond positively to the imitative acts of others extends to infants. When faced with an attentive adult, neonates imitate a limited range of the adult’s facial expressions (Meltzoff & Moore, 1977; Field, Woodson, Greenberg, & Cohen, 1982) and 3- to 5-month-old infants imitate a limited range of vocal expressions (Kuhl & Meltzoff, 1996), though the interpretation of these behaviors has raised controversy (see Ray & Heyes, 2010; Oostenbroek, et al., 2016). Imitation of both movements and vocal sounds becomes increasingly flexible and robust during the first year of life (Barr, Dowden & Hayne, 1996; Meltzoff, 1988; Jones, 2007). Shortly after their first birthdays, infants copy the behaviors of in-group members more closely than those of out-group individuals (Buttelmann, Zmyj, Daum & Carpenter, 2012; Howard, Henderson, Carrazza & Woodward, 2015). They also respond positively to being imitated, smiling more at an imitator than at a contingent social partner who does not imitate them, and helping more after being imitated by a friendly adult (Meltzoff, 1990; Agnetta & Rochat, 2004; Carpenter, Uebel & Tomasello, 2013). One-year-old infants also treat imitation as evidence of a robot’s capacity for social engagement (Meltzoff, Brooks, Shon & Rao, 2010). This research does not reveal, however, whether such infants possess an understanding of imitation that allows them to make the sorts of third party social inferences made by children and adults.

Studies of one-year-old children also do not shed light on the origins of an understanding of social imitation. The earliest imitative behavior may reflect only asocial sensory-motor associations; infants may come to endow imitation with social meaning by experiencing social interactions in which they are the initiator or the recipient of an imitative action (Cook, Bird, Catmur, Press & Heyes, 2014; Jones, 2006). Alternatively, the infant’s own imitative behavior may be social from the beginning, and supportive of social inferences (Meltzoff & Moore, 1992). Imitation is one of the very few communicative gestures used by adults that infants might, in principle, understand, because it does not require mastery of any culture- or language-specific conventions. Studies of young infants’ interpretations of imitation thus may shed light on infants’ social cognitive abilities more generally. We therefore investigate whether infants endow observed imitation with social meaning before they begin to engage in robust, socially motivated imitation of their own interactive partners.

Across these studies, we also ask whether the social information that infants gain from imitation applies symmetrically or asymmetrically to imitators and their targets. If infants view imitation simply as evidence that the target and imitator are similar, then they might make symmetrical social inferences about imitators and their targets. In contrast, if infants view imitation as a social behavior reflecting the imitator’s attention and/or motivation then infants’ inferences about imitators and their targets might differ. Although an imitator signals her social attention and motivation toward a target by her act of imitation, the target of imitation makes no such signal, if she does not respond to the imitator in turn.

### Current Studies

We report a series of experiments measuring the visual attention of 4- and 5-month-old infants who are presented with acts of imitation and social affiliation, together with one experiment conducted with 12-month-old infants. We use these patterns of attention to ask whether, after observing a series of imitative and non-imitative interactions, infants expect imitators and/or the targets of imitation to approach and affiliate with their partners in the imitative interaction. To convey imitation, we present characters who copy other characters’ movements or sounds. To convey affiliation, we use approach followed by synchronous motion. Approach is a basic behavior that is prompted by and indicates attraction to a person or object for adults (e.g. Cacioppo, Priester & Berntson, 1993; Chen & Bargh, 1999) and infants (Woodward, 1998; Gergely, Nádasdy, Csibra, & Bíró, 1995; Sommerville & Woodward, 2010; Martin, Vouloumanos & Onishi, 2012), and it has been used to test for expectations of positive social attitudes in infants (Kuhlmeier, Wynn & Bloom, 2003; Hamlin, Wynn & Bloom, 2007). Synchronized motion by animate characters also prompts social affiliation in adults (Hove & Risen, 2009) and infants (Cirelli, Einarson & Trainor, 2014), and it is interpreted by infants as a sign of social affiliation (Powell & Spelke, 2013). By testing for expectations of approach and/or synchronous motion, therefore, we ask whether infants infer that imitators possess positive attitudes toward the targets of their imitation and vice versa.

Experiments 1a and 1b tested whether infants expect an individual character who has imitated one pair of characters to affiliate with that pair relative to a different pair it did not imitate. Experiment 2a extended this question by testing whether infants respond similarly when, following the same imitative interactions, the pairs approach and affiliate with the individual instead. Experiment 2b reversed the roles of the individual and pairs in the imitative interactions (i.e. as imitators versus targets of imitation) and then tested separate groups of infants on trials in which either the individual or the pairs played the approaching role (see Figure 1). Experiment 3 tested for developmental changes in infants’ interpretation of these events, with an experiment similar to Experiments 1a and 1b.

Experiments 4 and 5 investigated the same questions in the context of dyadic interactions. In Experiments 2 and 4, moreover, some conditions presented test trials in which one actor approached two different parties, testing infants’ inferences concerning the approacher’s likely social goals or attitudes. Other conditions presented two different parties who both approached the same character, testing infants’ inferences regarding who will initiate the affiliative interaction.

The experiments have three notable features. First, they depict imitation and affiliative events using animated characters consisting of geometric shapes with faces that move spontaneously and produce sounds. Such characters readily elicit mental state and social inferences in adults, children and infants when they move in a self-propelled and goal-directed manner (Heider & Simmel, 1944; Johnson, Dweck & Chen, 2007; Kuhlmeier, et al., 2003; Over & Carpenter, 2009; Schachner & Carey, 2013; Hamlin et al., 2007; Thomsen, Frankenhuis, Ingold-Smith & Carey, 2011; Mascaro & Csibra, 2012; Powell & Spelke, 2013). Indeed, neurotypical individuals default to animate, social perceptions of simple shapes when presented with contingent, complex, and self-propelled behaviors (e.g. Heider & Simmel, 1944; Castelli, Happé, Frith & Frith, 2000). We thus chose novel, artificial sounds and movements that nevertheless are likely to be perceived by infants as voluntarily generated by animate entities (Powell & Spelke, 2013).

By presenting computer-animated stimuli as opposed to live, videotaped, or puppet-based displays, we can assure that all experimenters are naïve to the events seen by individual participants throughout the execution of the study. We also gain greater control over differences between conditions, varying only the patterns of imitation and approach between individuals and not other aspects of their actions. Finally, the use of animated characters allows us to present highly simplified events to young infants, whose processing of social events may be hampered by marked limits to their perceptual resolution and working memory.

A second feature of these experiments is that they compare infants’ inferences about the social behavior of groups vs. individuals. The first three experiments tested infants’ expectations concerning affiliative behavior following imitation conducted in a group context. We first focused on infants’ responses to imitation of or by groups because group contexts enhance social imitation for adults and young children, especially when the group is unanimous in its behavior or judgments (Haun, Rekers & Tomasello, 2012; Asch, 1956; Watson-Jones et al., 2016), and because there is evidence that infants expect social group members to act alike (Powell & Spelke, 2013). Providing a group context therefore might help infants to focus on the affiliative implications of imitation. After establishing the pattern of inferences infants make in this group context, the final two experiments investigated infants’ social interpretations of dyadic imitative interactions.

A third feature of these experiments is that they assume graded expectations regarding the likelihood of future social behavior. The most common prediction regarding infant visual attention is that infants will look longer at events or outcomes they find unexpected, compared to those that match their expectations. However, preverbal infants form graded expectations concerning the likelihood of different future events (e.g., Munakata, McClelland, Johnson & Siegler, 1997; Téglás, Vul, Girotto, Gonzalez & Tenenbaum, 2011), and these expectations do not map linearly onto the duration of their attention to those events (e.g., McCall & Kagan, 1967). Infants’ attention has been linked to an effort for efficient information gain, which is advanced by neither highly predictable (and thus uninformative) nor highly unpredictable (and thus uninterpretable) events. Instead, infants attend most to events of intermediate likelihood that support revisions of their predictions about or interpretation of a given context (Kidd, Piantadosi & Aslin, 2012).

The drive for information gain also explains variation in looking preferences for novelty versus familiarity. When a repeated display has been fully encoded, it offers little opportunity for further learning, and so infants tend to look longer at a novel display. When infants have not fully encoded a repeating display, in contrast, they have more to learn about it and may continue to look more at that display than at a novel one (McCall & Kagan, 1967; Turk-Browne, Scholl & Chun, 2008). Degree of encoding is affected by factors such as the amount of familiarization, the complexity of the events or entities depicted, and the maturational level of the cognitive processes recruited for encoding (Hunter & Ames, 1988; Roder, Bushnell & Sasseville, 2000).

Well-encoded displays may support the formation of strong expectations concerning future events. Under these conditions, infants are likely to view the expected test event as highly probable, and therefore show less interest in that event than in the unexpected event, which presents them with an opportunity to learn (Stahl & Feigenson, 2015). If the initial events are more complex, less compelling, or more demanding of attention and memory, in contrast, infants likely will form weaker expectations about future events. Under these conditions, test events that match the initial events may be more informative for infants, insofar as they strengthen the infants’ understanding of the original event. Thus, infants sometimes will attend more to events that are more expected.

Studies of older infants’ attention to imitation provide evidence for these graded expectations and diverging attentional preferences. At 8 months, infants reveal their expectation that a character will imitate the members of its own group by looking longer at an unexpected event (imitation of an outgroup) if the groups of characters are homogeneous in appearance, making group membership easy to encode and remember. In contrast, infants of the same age look longer at the expected event (ingroup imitation) when the groups are heterogeneous in appearance, making group membership more difficult to track. Testing older infants or scaffolding memory for group membership shifts attention back in the direction of unexpected events (Powell & Spelke, 2013). Studies of infants’ expectations of affiliation show a similar pattern. When presented with characters whose faces and behavior both mark them as clearly social, 10-month-old infants look longer at events in which a social character approaches another character who has previously hindered him, rather than a character who has previously helped him (Hamlin et al., 2007): they attend more to the unexpected test event. Conversely, when presented with characters that have no faces and whose animacy thus is more difficult to determine, 12-month-old infants look longer at approaches toward the helper (Kuhlmeier et al., 2003): they attend more to the expected test event.

When social events are ambiguous or difficult to encode and remember, therefore, infants form weaker social expectations and attend more to events that confirm those expectations. Given these findings, and our uncertainty concerning the complexity of the present experimental displays for young infants, we initially assessed expectations of approach following imitation by testing for visual attention to consistent and inconsistent events that differ in either direction (with an alpha value of p < 0.025 for each tail). We also varied the complexity of the displays, the amount of exposure to the displays that infants were given, and the age of the infants (4 vs. 12 months) to test whether these variables affect infants’ looking patterns in the ways that the hypothesis of graded expectations would predict.

## Experiments 1a and 1b

In Experiments 1a and 1b, we tested 4-month-old infants’ expectations concerning the affiliative behavior of a character who responds to the members of two different groups by imitating the distinctive action of one group of characters and not the other. In Experiment 1a, the responder imitated the sound made by one group and not the other, while movement was held constant across all characters. In Experiment 1b, the responder imitated the movement produced by one group and not the other, while sound was held constant across all characters. We tested infants’ expectations in the contexts of sound-based and movement-based imitation, because both are likely to be familiar and important imitative contexts for infants. Parents imitate infants’ vocalizations and their facial expressions (Kokkinaki & Kugiumutzakis, 2000; Pawlby, 1977), and past research suggests infants link both shared vocal behavior and shared movements to social affiliation (Kinzler, Dupoux & Spelke, 2007; Liberman, Woodward & Kinzler, 2016; Powell & Spelke, 2013). By testing infants’ responses in each context separately, we can assess the extent to which their responses to observed imitation generalize across these two modalities.

The two groups, each composed of two animated characters of the same color and standing in proximity, were present onscreen throughout the experiments, as was a fifth character of a different color, standing equidistant between the groups (hereafter, the “individual”) (Figure 1). Each familiarization event depicted the members of one group individually and sequentially moving the same way and making the same sound, followed by the individual also moving and making a sound. Participation in the events alternated back and forth between the two groups, and either the sounds (Experiment 1a, Figure 1a) or the movements (Experiment 1b, Figure 1b) made by the two groups differed. The individual consistently responded to *both* groups by producing the same sound and movement as one of the groups. Thus, in Experiment 1a, this character consistently responded to one group by imitating its sound, and consistently responded to the other group by making a different sound, while all the characters moved in the same manner. Conversely, in Experiment 1b, this character consistently responded to one group by imitating its motion, and responded to the other group by making a different motion, while all the characters emitted the same sound.^1^

**Figure 1.**
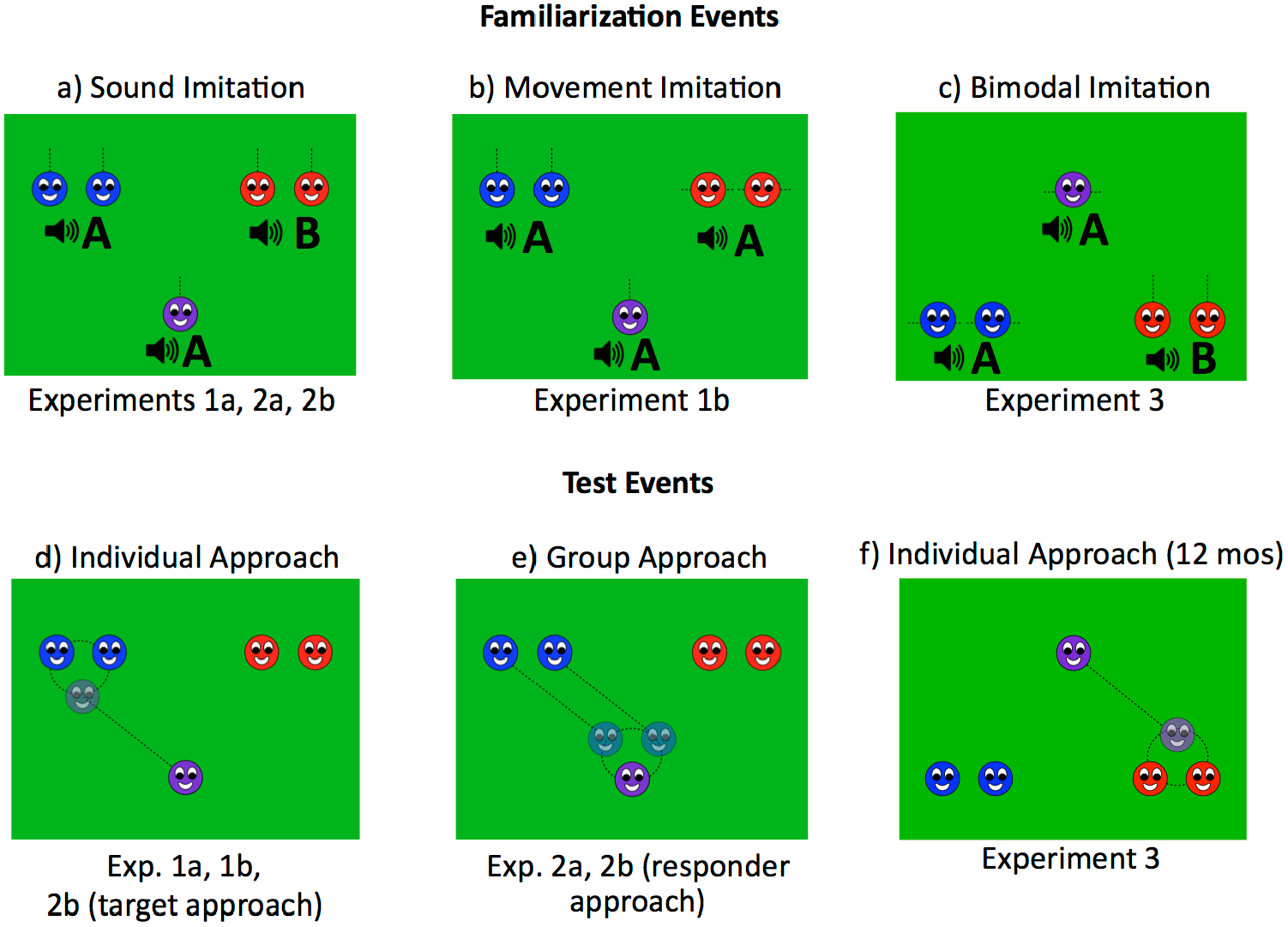
Example scenes from Experiments 1a, 1b, 2a, 2b, and 3. In familiarization events an individual character imitated either (a) the sound (Experiments 1a and 2), (b) the movement (Experiment 1b), or (c) both the sound and the movement (Experiment 3) made by one group but not the other. In test events either (d & f) the responding individual alternated in affiliating with the two groups (Experiments 1a, 1b, 2b, and 3), or (e) the initiating groups alternated in affiliating with the individual (Experiment 2a and 2b).

The familiarization events were interleaved with test trials in which the responding character approached and then moved synchronously with the members of one group, alternating between the groups across trials (Figure 1d). We contrasted infants’ looking times to test trials where the character approached and moved with each of the two groups, in order to assess the effect of imitation on infants’ expectations of such affiliative behavior.

### Experiment 1a

#### Methods

##### Participants

Participants were 16 4- to 5.5-month-old infants (8 female; age range: 4 months, 6 days – 5 months, 12 days; mean age: 4 months, 23 days). One additional participant was excluded for fussiness and one for experimental error.

##### Materials and Procedure

Participants sat in a car seat facing a 40 x 60cm display screen from a distance of approximately 60 cm. An inconspicuous camera above the infant recorded infants’ attention to the events; a camera to the side of the infant recorded the displays as they were presented. Infants watched animations (created and displayed using Keynote ‘08) featuring circular figures (each 6.5 cm in diameter) with schematic faces depicting eyes facing forward and a smiling mouth, against a uniform green background. There were two blue characters in the upper left corner of the screen, two red characters in the upper right corner of the screen, and one purple character centered toward the bottom of the screen (Figure 1a). The two characters within a group were separated by 4 cm, the two groups were separated by 23 cm, and the individual stood 28 cm from the midpoint of each group.

Participants viewed two rounds of events, each consisting of four familiarization events depicting imitative and non-imitative interactions followed by two test events depicting approach and synchronous movement. The familiarization events began with one member of a group jumping vertically three times and making the same sound at the initiation of each jump (Figure 1a). The second member of the same group then performed these actions, so that members of a group acted sequentially, first the left member and then the right member, with a pause of 1 s between the actions of the two characters. Following another 1 s pause, the responding character also jumped three times while making a sound. Participation in the familiarization events alternated between the blue and red groups, which each made different, group-specific sounds when jumping. The sound made by the responding character was constant and matched that produced by one of the groups but not the other. It thus comprised an imitative action when it followed one group and a non-imitative action when it followed the other group. These interactions each occurred twice in an alternating order. They were separated by 2 s pauses, during which a verbal cue from the experimenter (“Look, [baby’s name]!”) preceded each interaction.

The test events began with the experimenter calling to the baby (“Look, [baby’s name]!”) followed by a knocking sound, and then depicted the responding character approaching either the group it had imitated (a *congruent* event) or the group it had not imitated (an *incongruent* event) and then moving in synchrony with the two group members around a circular pathway, stopping after one full rotation (Figure 1d). Blind, online coding began at the point when the character met the group it was approaching and began moving with them, and continued until coding indicated that the participant had looked for 60 s cumulatively or had looked away for 2 s consecutively. These thresholds were set prior to the start of data collection. The two test events followed the same structure, each presenting affiliation between the responding character and a different group of characters. Once the look-away time threshold was met, the responding character moved back to its original position and the animation proceeded with the next event in the sequence.

##### Design

The order of familiarization to the imitative and non-imitative interactions, the group involved in the imitative interactions (red vs. blue), the sound that was imitated (high- vs. low-pitched), and the order of the test events (congruent or incongruent first) were all orthogonally counterbalanced between subjects. The animation was controlled and coded by display-blind experimenters, with coding initiated according to sounds associated with the characters’ actions.

##### Data analysis

Analyses were conducted on cumulative looking times following each test event. Looking times were recoded offline, also blind to condition, with 25% of participants coded by two independent experimenters. Measurements by the two coders were highly correlated (*r* = 0.99). To weight the relative difference between looking times to congruent and incongruent events equally across the two test pairs, we calculated the proportion of the looking time for each pair of test events that was spent looking to the incongruent event and then averaged this proportion across the two pairs of events. If an infant did not watch an event for at least 0.5 s or if some source of experimental error occurred during an event (e.g. flawed online coding resulting in a trial cut short; parental interference), then the pair of events to which this trial belonged was excluded (a total of 4 trial pairs from 4 different participants were excluded on this basis). The proportion of looking to the congruent test events for each infant was compared to chance (50%) by a one-sample, two tailed t-test. A single ANOVA tested for effects of or interactions between familiarization order (imitation or non-imitation first) and test order (congruent first or second) as well as effects of gender on looking time to the congruent trials.^2^ A similar single ANOVA testing for familiarization and test order effects was performed in subsequent experiments as well, though the factor of gender was dropped due to its failure to show any evidence of interacting with looking times.

#### Results

Infants looked proportionally longer at congruent test trials, in which the responding character approached and moved with the group of characters it previously imitated (59.4%, *t*(15) = 3.09, *P* < 0.01; Fig. 2a). The ANOVA found no effects of event order or gender on this difference. The looking preference provides evidence that infants distinguished the congruent from the incongruent approach trials, and were more motivated to attend following congruent trials. Before pursuing the source of this motivation, we tested whether the same effects would be obtained when infants viewed imitation of motions rather than sounds.

### Experiment 1b

#### Methods

##### Participants

Participants were 16 4- to 5.5-month-old infants (11 female; age range: 4 months, 3 days – 5 months, 14 days; mean age: 4 months, 17 days). One additional participant was excluded due to experimental error.

##### Materials and Procedures

The method was the same as in Experiment 1a, except as follows. During the familiarization events, the members of one group jumped vertically three times exactly as in Experiment 1a, making the lower-pitched sound from that experiment. The members of the other group made the same sound but a different movement: they slid horizontally three times for a distance of 13 cm, with the sound synchronized with the endpoints of the motion. The responding character performed one of these two actions (accompanied by the same sound), thus imitating the movement of one group and not the other (see Figure 1b). The primary data analyses were the same as in Experiment 1a. Five participants had one pair of trials excluded due to the ineligibility of one or more trials in the pair. All participants were recoded by one or more blind, offline coders, and correlation of looking times recorded by two independent coders was high (*r* = 0.99 across 25% of participants). Finally, an independent sample t-test compared infants’ proportional looking to congruent trials across the two experiments. When no differences were found, a one sample t-test on the combined findings from the two experiments served to estimate the size of the congruency preference effect.

**Figure 2.**
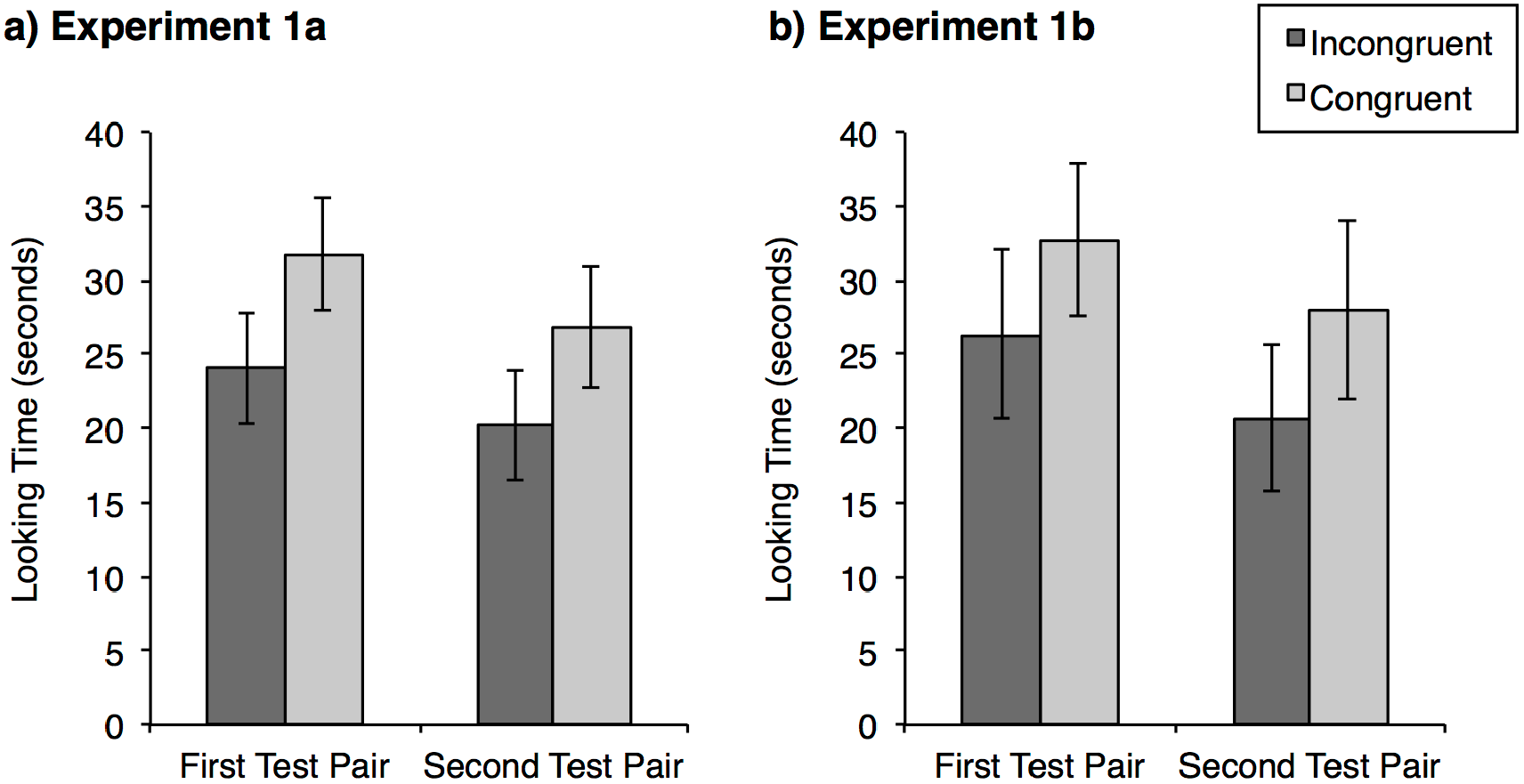
Mean looking times recorded during the two test pairs for (a) Experiment 1a and (b) Experiment 1b. Error bars represent standard error of the mean (SEM).

#### Results

Infants again spent more of their total looking time attending to the congruent trials, in which the individual approached and danced with the group it imitated (*M* = 58.5%, *t*(15) = 2.66, *P* < 0.05; Fig. 2b). The ANOVA testing for effects of familiarization order, test order and gender revealed no significant main effects or interactions. The proportion of looking to congruent test trials did not differ across the two experiments (*t*(30) = 0.02, *P* > 0.8), which together showed a strong effect of congruency with prior patterns of imitation on infants’ looking to affiliative approach events (*t*(31) = 4.12, *P* < 0.001, Cohen’s *d* = 0.73).

### Discussion

In Experiments 1a and 1b, infants consistently looked longer to events in which an individual approached and moved in concert with the group it had imitated, compared to those in which the individual approached and interacted with the non-imitated group. One explanation for this finding is that it reflects a low-confidence expectation of affiliation between the individual and the imitated group, rather than between the individual and the group not imitated. On this interpretation, events in which the individual approached the imitated group provided confirmatory evidence that the individual was, in fact, positively oriented toward the imitated group, whereas the incongruent approaches failed to contribute to a coherent understanding of the characters and events. Thus infants may have devoted more attention to the congruent, confirmatory events because they found them more informative. On this account, infants’ converging responses to imitation of sounds and movements is consistent both with the nature of the social imitation in infants’ environments and with the responses of children and adults, for whom vocal and motor mimicry both reflect and elicit positive regard (Giles & Powesland, 1975; Neumann & Strack, 2000; Adank et al., 2010; Chartrand & Lakin, 2013).

However, a weak expectation of affiliation between an imitator and its target(s) is only one of many potential accounts for the present findings. Several leaner explanations for these findings are possible. In particular, infants may not construe the familiarization events in terms of imitation (i.e. as the responding character repeating the behavior of the two characters in one of the two groups but not the other). Instead, infants may perceive only that the sound or motion produced by the individual is more similar to one group than to the other. This observation alone could provide grounds for infants to distinguish the congruent and incongruent test trials. The sameness and ease of processing of the homogenous group that resulted from the congruent approach, or a preference based on the accumulated familiarity of the imitated actions, more often repeated than those of the non-imitated group, may have elicited greater attention from infants on those trials (Hunter & Ames, 1988; Zajonc, 1968).

Even assuming the events in Experiment 1 were perceived as social interactions, several alternative sources of an expectation that the individual would affiliate with the imitated group should be considered. First, infants may have inferred some likelihood that the individual and the imitated group were affiliated, but on the basis of a principle of homophily rather than an understanding of imitation. Past research demonstrated that infants expect members of social groups to act alike (Powell & Spelke, 2013); infants’ longer looking to the congruent events in Experiments 1a and 1b may be driven by a tentative inference that individuals who act alike are part of the same social group. Second, the imitation may have increased infants’ impression of the imitated group’s importance or likability. If so, then the preference for congruent trials could reflect a weak expectation that *any* individual would be more likely to approach the imitated group.

Finally, infants’ preference to attend to the congruent events may not reflect the expectedness of the affiliation events at all, but rather infants’ own social preference or their anticipation of events to come. If infants do perceive the imitated group as more desirable, congruent test events may have elicited longer looks because they drew infants’ attention toward the more desirable target. Or, the increased looking could have been driven by an anticipatory expectation that the individual and the imitated group would continue to interact with one another.

The next experiments aimed to test the hypothesis that infants view imitation as a reflection of the imitator’s social attention and motivation, against all these alternatives, by comparing infants’ expectations about the affiliative behaviors of imitators with their expectations about the affiliative behavior of the targets of imitation. We continued to present infants with events that followed the same basic structure while varying the roles of the characters in the imitation and approach events as well as the number of characters onscreen, so as to test infants’ expectations of groups as well as individuals, and of targets of imitation as well as imitators. These variations allowed us to isolate the features of Experiments 1a and 1b that are critical for infants’ differentiation of imitation-congruent and –incongruent approach and interaction. Together, they also allow us to test further whether the direction of infants’ looking preferences indeed was modulated by the complexity of the displays.

## Experiment 2a and 2b

Experiments 2a and 2b compared infants’ expectations about the affiliative behavior of imitators, as in Experiment 1, to their expectations about the affiliative behavior of targets of imitation. To this end, we manipulated the roles of the characters in both the imitative interactions and the subsequent affiliation events. Experiment 2a presented infants with the same imitative events as in Experiment 1a: two pairs of characters each jumped while making a distinctive sound and an individual character responded to each pair by making one of the two sounds, thereby imitating one group of characters and not the other. For the test trials, however, we exchanged the roles played by the groups and the individual. Rather than presenting the individual alternately approaching the two groups, infants were presented with events in which the two groups alternately approached and moved synchronously with the individual (Figure 1e). This change also reversed the relationship between the imitative and affiliative roles: the affiliation events were now initiated not by the imitator but by the targets and non-targets of imitation. We tested whether infants again show greater attention to the congruent test event displaying affiliation by the group that was the target of imitation.

We also tested infants in two further conditions (Experiment 2b) in which we reversed the order of actions in the imitation events, such that the lone character acted first and the characters in the two groups responded to its action, one group imitating the action and the other group performing a different action. These events were followed, for different groups of infants, by the two different types of affiliation events presented in Experiments 1a and 2a (i.e. individual approach or group approach). Due to the inversion of the imitation events, the test events depicting the individual approaching each of the groups now portrayed the *target* of imitative and non-imitative actions approaching the authors of those actions (and are thus referred to as the Target Approach condition), while the events depicting the groups approaching the individual represented responding parties approaching the target of their imitative or non-imitative acts (Responder Approach condition).

Together with Experiment 1a, the three conditions in Experiments 2a and 2b complete a 2 x 2 design in which the two versions of the imitative interactions, the one in which the groups initiate and the individual responds and the other with the reverse order of actions, are each separately paired with affiliation events in which the individual approaches the groups and ones in which the groups approach the individual. Comparing infants’ patterns of attention to the affiliation events across these four conditions allows us to test whether greater attention to congruent than incongruent trials was observed universally, was dependent on particular imitation or affiliation displays, or was linked to the relationship between the two types of displays.

### Experiment 2a

#### Methods

##### Participants

Participants were 16 4- to 5.5-month-old infants (8 female; age range: 4 months, 2 days – 5 months, 14 days; mean age: 4 months, 21 days).

##### Materials and procedure

The procedure, design, dependent measures and data analysis were the same as in Experiments 1a and 1b. The displays were the same as in Experiment 1a except that, rather than seeing the individual approach each of the groups during the test events, the groups now alternately approached the individual. These approach events were followed by the same synchronized, circular movement as in Experiment 1, but the movement now took place near the original position of the individual character (Figure 1e). The test events presenting approach by the imitated and non-imitated groups were considered congruent and incongruent, respectively. Two pairs of trials, one each from two different participants, were excluded due to the ineligibility of one or more trial from the pair. All looking times were recoded by one or more blind, offline coders, and correlation of looking times recorded by two independent coders was high (*r* = 0.99 across 25% of participants).

#### Results

Infants looked no more at the congruent test events (47.6%) than at the incongruent events (*t*(15) = 0.65, *P* > 0.5, Figure 3b). The two-way ANOVA testing for effects of familiarization and test order also revealed no main effects or interactions (all *P* > 0.5). This lack of differentiation between the test trials represented a substantially different pattern of attention than that observed in Experiments 1a (Figure 3a) and 1b. An independent samples t-test comparing proportion of looking time to congruent trials confirmed this difference (*t*(46) = 2.83, *P* < 0.01).

### Experiment 2b

#### Methods

##### Participants

Participants were 32 4- to 5.5-month-old infants (16 female; age range: 4 months, 1 day – 5 months, 15 days; mean age: 4 months, 24 days). Two additional participants were excluded for fussiness.

##### Materials and Procedure

The method was the same as that used in Experiments 1a and 2a, except as follows. Infants were pseudorandomly assigned to a Responder Approach or Target Approach condition, equating for gender across the two conditions. For both conditions, the order of characters’ participation in the familiarization events was reversed. These events began with the individual character initiating an interaction by jumping and making the same sound three times, followed by the members of one group responding in sequence by jumping and making a sound as well. Both members of one group made the same sound as the individual character, such that the group now imitated the individual; both members of the other group made a different sound and therefore did not imitate the individual target character.

**Figure 3.**
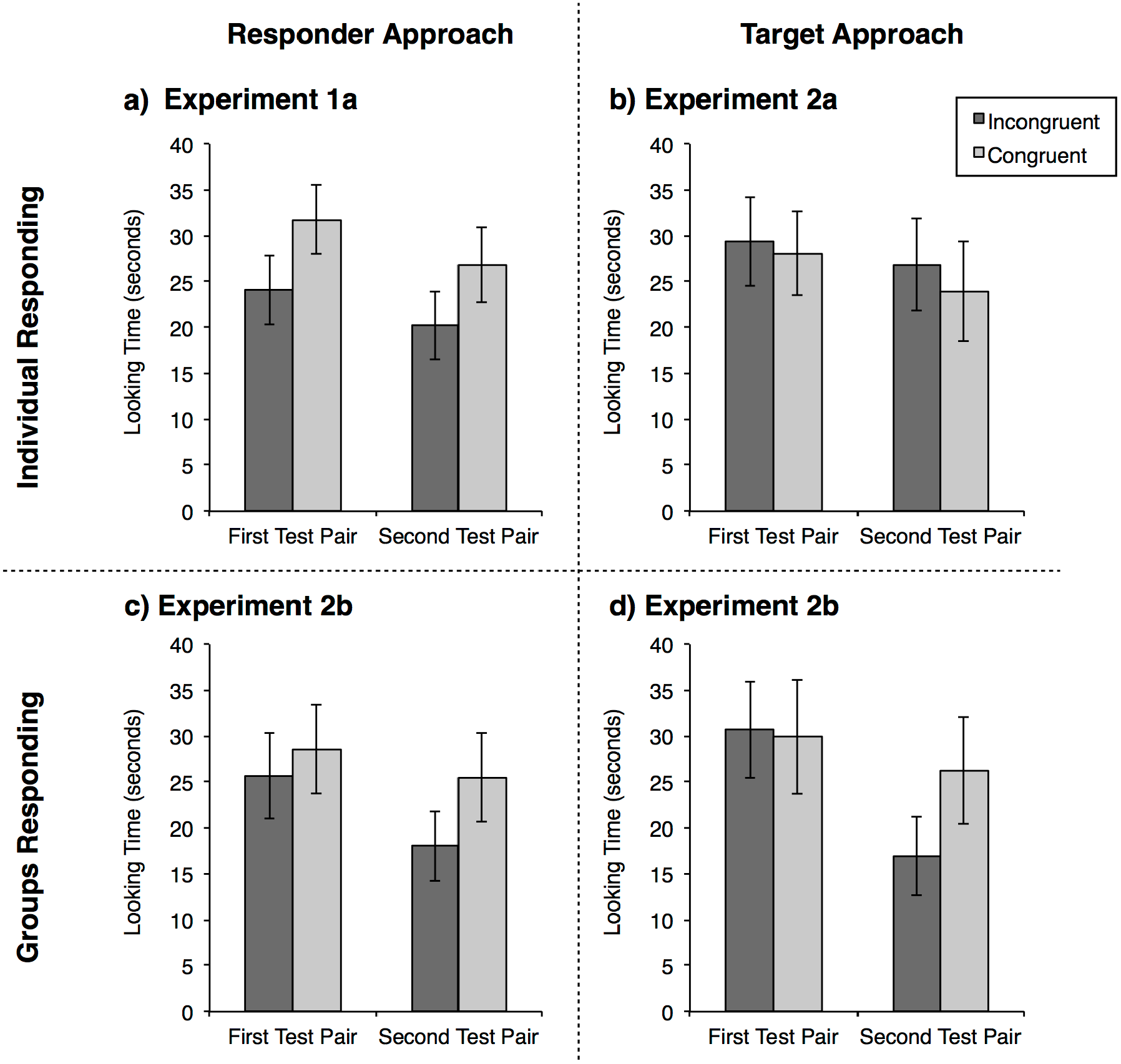
Mean looking times by test pair for Experiments 1a, 2a, and 2b, representing a 2 x 2 matrix of sound imitation events that varied the identities of the responders and targets (lone character as responder, grouped characters as targets represented in top row; reversed identities represented in bottom row), as well as the role played by the approacher in the interaction events (responder approach represented in the left column, target approach in the right column). Error bars represent SEM.

The test events of the Responder Approach condition were the same as those of Experiment 2a: the two groups alternately approached and moved in synchrony with the initiating character. This condition therefore assessed infants’ attention to events in which a lone individual is approached, in alternation, by groups that previously imitated or failed to imitate its actions. The test events of the Target Approach condition were the same as those of Experiment 1a: the lone character alternately approached and moved in synchrony with each of the two groups. This condition therefore assessed infants’ attention to events in which a lone individual alternately approached groups that previously imitated or failed to imitate its actions. One pair of trials each was excluded for six different participants. All looking times were recoded by a blind, offline coder; correlation between looking times recorded by the two independent coders was high (*r* = 0.98 across 25% of participants).

##### Data Analysis

As in the previous experiments, we compared the proportion of looking directed to congruent trials to chance (50%) for each condition. To assess whether this proportion differed across the Target Approach and Responder Approach conditions, condition assignment was included as a between-subjects factor in an ANOVA also testing for effects of imitation order and test order.

After investigating the pattern of looking behavior within Experiment 2b, we compared looking times across Experiments 1a, 2a and 2b, encompassing the 2 x 2 comparison within which the factor of individual response to the groups or group responses to the individual during familiarization was crossed with the factor of individual approach or group approach at test. We conducted an ANOVA examining proportion of looking time to congruent trials, with responder and approacher identity (lone character or group for each variable) as between subjects factors, along with test order, as this factor was observed to have an influence in Experiment 2b. If specific familiarization or test displays drove – or masked – infants’ differentiation of congruent and incongruent approaches, they should produce main effects of the responder or approacher factors. If, instead, infants show a congruency bias when an imitator approaches a target but not the reverse, then across all four conditions we should see a responder x approacher interaction, reflecting a crossover pattern in which infants devote more looking time to congruent trials when test events depict approach by imitators rather than by targets.

#### Results

Infants in the Target Approach condition looked equally at the congruent (50.5%) and incongruent test events (*t*(15) = 0.11, *P* > 0.9; Figure 3d). Participants in the Responder Approach condition also showed no significant looking preference between the test events (*M* looking to congruent = 55.7%, *t*(15) = 1.85, *P* = 0.08; Figure 3c), but there was a non-significant trend toward longer looking at the congruent test event. The ANOVA testing for effects of condition, imitation order and test order across the two groups found no main effect of condition but a significant interaction between condition and test order: Infants in the Responder Approach condition who saw congruent trials first looked longer to the congruent trials (63.9%, *t*(7) = 4.36, *P* < 0.01) and longer than infants who saw the incongruent trials first (*t*(14) = 3.53, *P* < 0.01). The latter group’s attention to the congruent trials did not differ from chance (47.5%, *t*(7) = 0.48, *P* > 0.7), nor did looking for infants in the Target Approach condition who saw congruent trials either first (48.8%, *t*(7) = 0.17, *P* > 0.8) or second (52.1%, *t*(7) = 0.42, *P* > 0.6). Though these results suggest that, for infants in the Responder Approach condition, a greater interest in earlier test trials competed with a congruency bias, the small sample size and the absence of a test trial order effect in the Target Approach condition, or any of the preceding experiments, make it difficult to draw conclusions from this effect.

The findings of the ANOVA comparing the proportion of looking devoted to congruent test trials across Experiments 1a, 2a and 2b, were more clear. Neither the identity of the responder in the familiarization events nor the identity of the approacher in the test events had an independent effect on infants’ relative attention to congruent events (both *P* > 0.3), but these two factors showed a significant interaction (*F*(1,56) = 6.01, *P* < 0.05). Infants devoted greater attention to the congruent than the incongruent test events when the identity of the responders and approachers matched across the two event types (i.e. the individual both responded and then approached or the groups both responded and then approached: *M* = 57.9%, *t*(31) = 3.51, *P* < 0.005; Figure 3a and 3c), but did not look longer at the congruent events when the responding party and the approaching party differed (*M* = 49.0%, *t*(31) = 0.35, *P* > 0.7; Figure 3b and 3d). Thus infants’ differentiation of the test events depended not on any particular feature of the approach test displays or the preceding interaction events, but rather on the relationship between the roles the characters played, as imitator vs. target of imitation, across the two types of events.

#### Discussion

Experiments 1 and 2 reveal a consistent asymmetry in infants’ responses to the affiliative behavior of imitators and their targets. Infants reliably differentiated events in which a responding party, whether a lone individual (Experiment 1; Figure 2 and Figure 3a) or a group composed of two individuals (Experiment 2b, Responder Approach condition; Figure 3c), approached and interacted with a target party it had imitated, from events in which the same or another responding party approached a target party it had not imitated. In contrast, infants failed to differentiate cases in which the initiators of interactions approached imitating versus non-imitating responders, regardless of whether the initiators were two groups (Experiment 2a; Figure 3b) or a lone character (Experiment 2b, Target Approach condition; Figure 3d). This asymmetry is particularly striking, because exactly the same familiarization and test events were alternately used to create both responder approach and target approach conditions across the four experiments. Infants responded to the pairing of these displays, not to the features of any particular display.

The cumulative analysis of these four experiments speaks against a number of potential low-level explanations for infants’ looking behavior. In particular, infants’ looking preference for congruent approach trials in responder approach conditions cannot be explained by a general preference for characters whose behavior is more familiar. Although the three characters involved in the congruent events of the responder approach conditions all made the movements or sounds that were imitated and were thus presented more frequently than the non-imitated movements or sounds, the same was true for the characters involved in the congruent events of the target approach conditions, in which no looking preferences were observed.

The results also rule out the possibility of a general preference to look at more homogenous collections of entities, because the test events in different experiments presented identical degrees of similarity within and spacing between groups, yet elicited different looking behavior depending on the order of characters’ participation in the imitative actions that preceded these events. More generally, infants’ expectations regarding approach were based not on inherent properties of the characters or of the displays but on the roles that characters played in the preceding social interactions, consistent with a social analysis of the elements of contingent interaction, approach, and synchronous motion used to compose the displays.

The present findings also constrain hypotheses concerning the nature of this social analysis. In particular, they provide evidence that infants do not expect affiliation on the basis of the similarity among individuals alone, due to a third party expectation of homophily. They also do not expect affiliation between characters who share behaviors, as one would expect if shared behaviors were interpreted as a marker of membership in a shared social group or of adherence to shared social norms. If infants expected social characters to affiliate selectively with others who are similar to themselves, or with others who adhere to the same social norms, then infants should make symmetrical predictions of affiliation by imitators and their targets. Instead, infants perceived the order in which the characters acted during the imitative and non-imitative familiarization trials, and this order played a role in their generation of expectations about further efforts toward affiliation.

A finding that young infants make asymmetric inferences about imitators and their targets would have important implications both for theories of the nature and development of imitation, and for theories of the development of social cognition more generally. Before we consider those implications, however, an outstanding question must be addressed. Experiments 1 and 2 find that infants look relatively longer to cases of approach by responders toward imitated interaction partners compared to non-imitated ones: the test events that are congruent with their putative expectations. In most experiments, however, infants reveal their expectations by looking longer at events that are incongruent with their expectations: congruency preferences only are found when infants form weak or low-confidence expectations, leading to longer looking at the confirmatory test events.

Did the 4-month-old infants in Experiments 1 and 2 form weak expectations that imitators were disposed toward affiliation with their targets? Experiments depicting the actions of five different characters may place high demands on young infants’ attention and memory (Wood, 2007), reducing their confidence in the predictions that result from those actions and leading them to seek confirmatory evidence (Kidd et al., 2012; Kinney & Kagan, 1976; Hunter & Ames, 1988). In past research presenting similar imitative interactions, however, older infants have successfully navigated such demands, and have shown the signature preference for incongruent social actions involving as many as six distinct characters (Powell & Spelke, 2013). To test further our interpretation of the findings of Experiments 1 and 2, therefore, the next experiment presented events similar to those of Experiments 1a and 1b to a group of 12-month-old infants. If infants expect imitators to affiliate with their targets both at four months and beyond, and if younger infants’ longer looking at events that are congruent with this expectation reflects their uncertainty about what has taken place, due to the high demands of these events on their attention and memory, then the direction of looking preferences should reverse between the younger and the older age. Like the older infants in studies of infants’ expectations of imitative behavior (Powell & Spelke, 2013), 12-month-old infants should look longer at the incongruent test events.

## Experiment 3

In this experiment, 12-month-old infants were presented with events involving the same five characters, actions and sounds as Experiment 1, but the characters appeared in a different spatial arrangement and the imitative interactions of Experiments 1a and 1b were combined, such that the two groups were distinguished *both* by their motion and by their sound.^3^ If infants have graded expectations that imitators will approach their targets, and these expectations were weak in Experiment 1 and 2 because 4-month-old infants were overtaxed by the task of remembering and reasoning about the imitator-target relationships, then the 12-month-old infants in Experiment 3 should have stronger expectations of approach, and should reveal these expectations by looking longer at the incongruent, rather than the congruent, test event.

### Methods

#### Participants

Participants were 16 11.5- to 12.5-month-old infants (6 female; age range: 11 months, 17 days – 12 months, 10 days; mean age: 12 months, 0 days). Two additional infants were excluded, one as a result of technical failure, and one due to fussiness.

#### Materials and procedure

The procedure, design, dependent measures and data analysis were the same as in Experiments 1a and 1b, but the displays differed from those of Experiment 1 in several respects. The characters did not begin on screen, but rather entered in an introduction sequence in which (1) one set of paired characters entered from the left side of the screen, moved in a synchrony around a circular path, and then came to rest in their typical position, (2) the other pair did the same, after entering from the right side, and (3) the lone character entered from the top of the screen. Then the familiarization events commenced. To accommodate the lower tolerance of older infants for long preferential looking experiments, the movements of the different group characters were overlapped in time. The second character in the group began to move 0.2 s after the first character such that their actions seemed coordinated but not perfectly synchronized. A single set of sounds accompanied the movements, timed with the onset of the first character’s movements. The sequence of familiarization events was also altered such that infants first saw both groups act, without an intervening action by the responder following the first group, and then saw the responder choose which of the two groups to imitate. The spatial arrangement of the characters was inverted, such that the grouped characters appeared on the bottom of the display and the responding character was centered above them. Finally, as noted above, the two groups differed both in their sounds and in their motions. Each of the sounds from Experiment 1a was paired with one of the motions from 1b. These pairings were constant, but the action profile imitated by the responding character was counterbalanced across participants (Figure 1c). Three pairs of trials, one each from three different participants, were excluded due to the ineligibility of one or more trials from the pair. All looking times were recoded by blind, offline coders, and correlation of looking times recorded by two independent coders was high (*r* = 0.99 across 25% of participants).

#### Data Analysis

There was no factor of familiarization order because the imitator no longer had two separate interactions with the target and non-target groups, so to test for test order effects, we did an independent sample t test on the proportion of incongruent looking by infants who saw incongruent trials first versus second. We also conducted an independent samples t test comparing proportion of looking to incongruent trials in this experiment to that in Experiments 1a and 1b, conducted with similar displays but substantially younger participants. Because we were testing the hypothesis that older infants would be better able to process the displays and generate more robust expectations about the imitator’s social motivation, we tested the one-tailed hypotheses that proportion of looking to *incongruent* trials would be greater than chance and greater in Experiment 3 than in Experiment 1.

### Results

Infants looked significantly more at the *in*congruent test events (57.3%) than at the congruent test events (*t*(15) = 2.01, *P* < 0.05, Figure 4a). The independent samples t test comparing looking for the two test orders was not significant (*P* > 0.4). The t test comparing the current experiment to Experiments 1a and1b (Figure 4b), found that older infants devoted a significantly greater proportion of looking to incongruent trials than younger infants did (*t*(46) = 4.06, *P* < 0.0001). Thus, 12-month-old infants showed the opposite looking pattern to their 4-month-old counterparts in Experiments 1a and 1b.

### Discussion

In Experiment 3, 12-month-old infants looked longer at the test event in which the responding character approached the group that it had *not* imitated: the more common looking pattern in violation-of-expectancy experiments, providing evidence that the infants expect a lone character who imitates one of two groups of characters to affiliate with the group that it imitated. These findings provide an instructive contrast to the findings of Experiment 1, conducted with four-month-old infants. Although the infants at the two ages viewed the same types of characters, actions, and group-specific responses by the responding character, they showed opposite looking preferences to these events. The reversal, with increasing age, from a preference for the familiar to a preference for the novel event is consistent with the thesis that infants at both ages are sensitive to imitation and expect imitator characters to approach their targets, but that the present experiments place high demands on young infants’ attention and memory.

**Figure 4.**
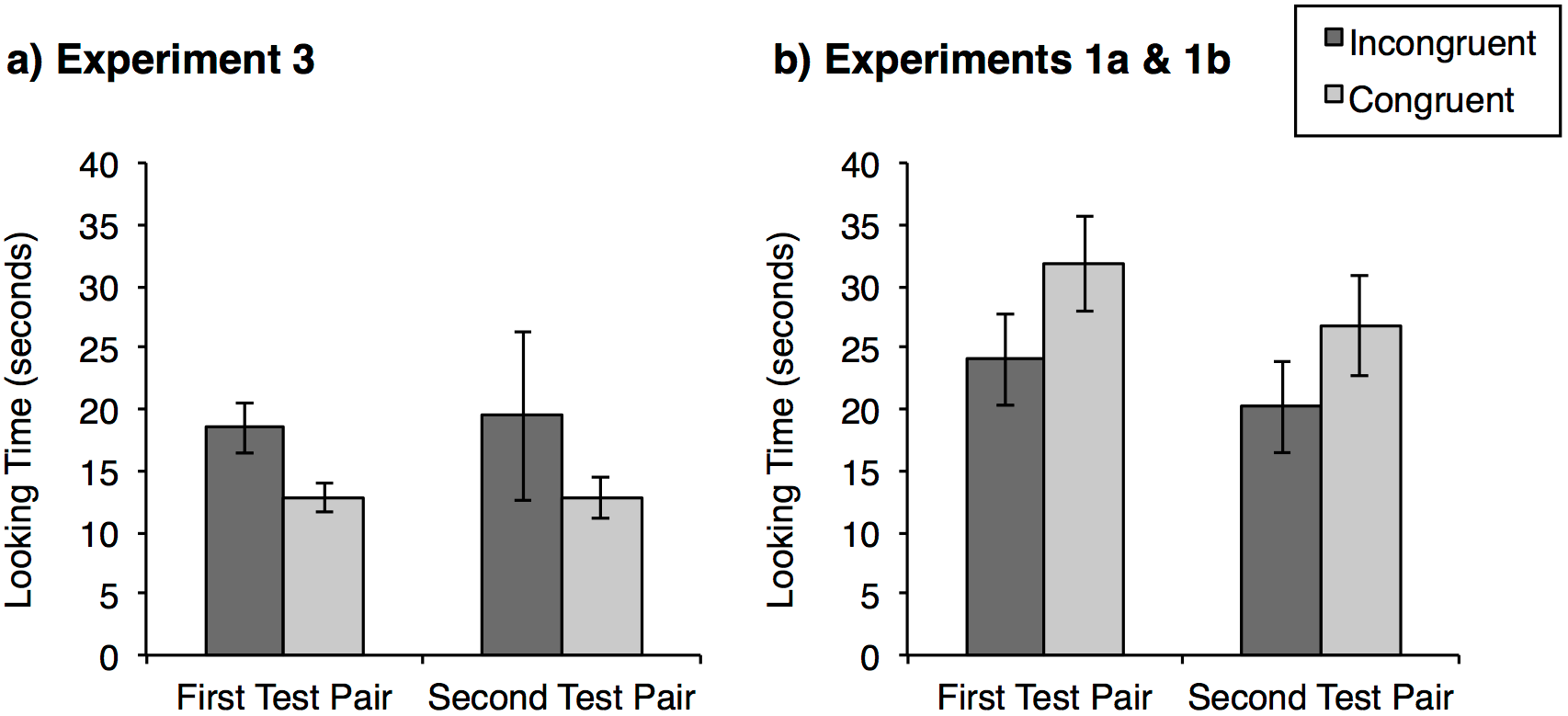
Mean looking times for each test pair in (a) Experiment 3, conducted with 12-month-old infants, and (b) combined for both Experiments 1a and 1b, conducted with 4- and 5-month-old infants. In contrast to data collected from younger infants, older infants spent more time looking to incongruent than to congruent trials. Error bars represent SEM.

This thesis can be questioned, because the events presented in Experiment 3 differed from those in Experiments 1 and 2 in a number of respects: they presented two socially distinctive actions rather than one, faster events, and a different spatial arrangement of characters. Might these changes, rather than the difference in age, account for the reversal in infants’ looking preferences from 4 to 12 months?^4^ Because the events did not differ from those of Experiment 1 in any way that affects the impression of imitative and affiliative behavior for adult observers, we believe that Experiment 3 served as a conservative test of the prediction that both infants at both 4 and 12 months expect the responding character to affiliate with the target group that it imitated, but that the direction of their looking preferences will reverse. Nevertheless, experiments that build more directly on the methods of Experiments 1 and 2 with participants of the same age, while reducing some of those experiments’ cognitive demands, would provide additional evidence that the results of those experiments reflected younger infants’ graded expectations.

If the events of Experiments 1 and 2 were indeed highly challenging for young infants, who formed weak expectations that imitators would affiliate with their targets, then a different possible interpretation of the imitator-target asymmetry found in Experiments 1 and 2 should be considered. Young infants’ asymmetric expectations regarding the affiliative behaviors of imitators and their targets, may stem from the added task demands of reasoning about targets of imitation, rather than from any asymmetric expectations about the behavior of imitators and their targets. If infants have only fragile expectations of affiliation by imitators, due to the high demands on attention and memory posed by the present events, then the asymmetry between young infants’ expectations about imitators and their targets might reflect the even higher demands posed by the test events in which the initiator(s) of the imitative interactions approached their responder(s). In those conditions, infants must use the imitative behavior of one character or group (the responder) to make an inference about the affiliative behavior of a different character or group (the target). Such an inference may be too difficult for infants when memory demands are high, resulting in a failure to differentiate the trial types even if infants are, in principle, capable of making inferences regarding the likely social affiliation of targets toward those who imitate them.

Two more experiments were undertaken to distinguish between the different interpretations of infants’ congruity preferences, and to test further infants’ expectations about the targets of imitation. In Experiments 4 and 5, we reduced the cognitive demands on infants by presenting three rather than five characters, replacing the pairs of red and blue characters with a single character of each color. We also decided in advance to increase the sample size (from 16 infants to 24 infants per condition) to increase the sensitivity of our tests. Experiment 4 consisted of four conditions analogous to those of Experiments 1 and 2. To test the effects of display complexity and memory demands on infants’ looking patterns, we compared infants’ responses to the new three-character events to their responses to the corresponding five-character events. In Experiment 5, we singled out the simplest condition from Experiment 4, in which an individual imitated one lone social partner but not another and then alternately approached each of these two targets at test. We replicated this condition with a change in method aimed at strengthening infants’ memory for the imitative interactions.

## Experiment 4

The four conditions in Experiment 4 repeated the four sound-based imitation experiments presented above with three characters rather than five (Figure 5). Two of the conditions, analogous to Experiments 1a and the responder approach condition of Experiment 2b, presented responder approach test trials in which the approaching character(s) alternately moved toward targets they imitated in prior interactions (congruent events) and those they did not imitate (incongruent events). If infants looked longer at congruent responder approach events in the five-character experiments because they expected imitators to approach their targets but high demands on attention and memory reduced their confidence in these predictions and increased their interest in confirmatory events, then reducing the number of characters to be tracked should strengthen infants’ expectation thereby weakening congruency preferences and increasing interest in the incongruent trials. In contrast, if infants looked longer at the congruent responder approach events because they were more intriguing or attractive for some other reason, then the congruency preference should be as strong or stronger in Experiment 4 as in the five-character experiments.

The remaining two conditions in Experiment 4 presented target approach conditions analogous to Experiment 2a and the target approach condition of Experiment 2b. If infants make asymmetric predictions about imitators and their targets, predicting social approach by imitators but not by targets of imitation, then infants should show the same absence of looking preferences in these three-character studies as in their predecessors. In contrast, if infants expect targets to approach their imitators, but failed to exhibit this expectation in previous experiments due to the high cognitive demands posed both by the use of five characters and the task of inferring the action of one party (the target) based on the actions of another party (the imitator), then the easing of memory demands in Experiment 4 may yield positive findings.

**Figure 5.**
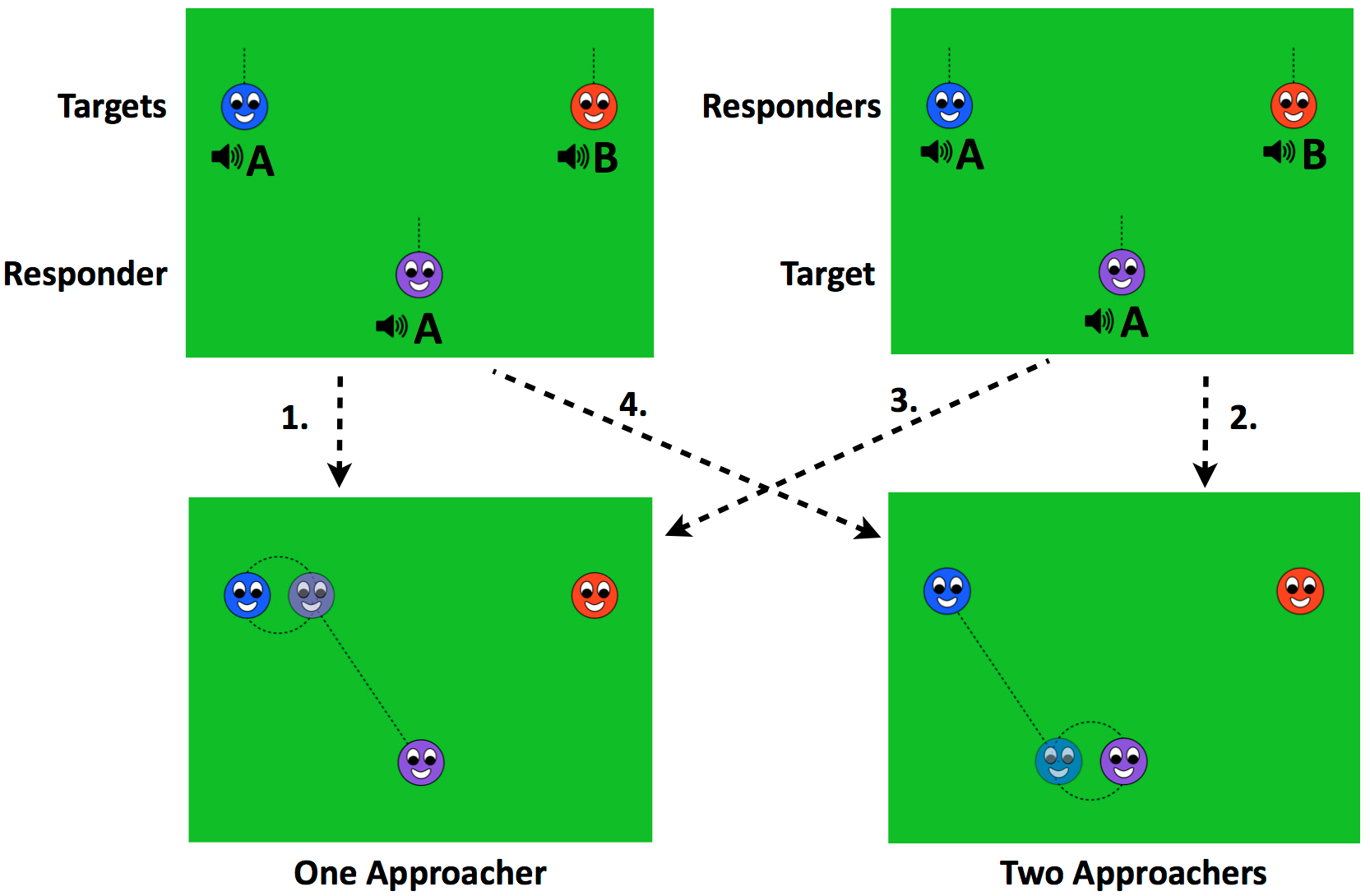
Example scenes from Experiment 4. Two types of familiarization events were crossed with two types of test trials to create four conditions. (1) Responder Approach, One Actor: Central character responded to each side character, then alternately affiliated with each. (These familiarization and test events were also used in Experiment 5.) (2) Responder Approach, Two Actors: Side characters each responded to central character, then alternately affiliated with central character. (3) Target Approach, One Actor: Side characters each responded to central character, then central character alternately affiliated with each. (4) Target Approach, Two Actors: Central character responded to each side character, then side characters alternately affiliated with central character.

The four conditions in Experiment 4 also present two different contrasts between test trials. In one responder approach and one target approach condition, the same actor (the central character) alternately approaches the other two characters (hereafter, "one-actor" conditions). Looking times measured across these congruent and incongruent trials thus assess infants’ relative expectations regarding the affiliative action that a single individual will take. In the other responder and target approach conditions, two different actors (the side characters) alternately approach the central character ("two-actor" conditions). The congruent and incongruent trials therefore assess infants’ relative expectations concerning who will undertake a given action. One-actor conditions may present a simpler inference problem for infants. Under those circumstances, the relative difference in the actor’s attitude toward its two partners can be inferred directly from its own, or its partner’s, past behavior. In two-actor conditions, in contrast, infants must infer how different interactions reflected the relative attitudes of two separate parties. The four conditions of Experiment 4 make it possible to assess the impact of this factor on expectations in both responder approach and target approach contexts.^5^

### Methods

#### Participants

Participants were 96 4- to 5.5-month-old infants (24 in each of 4 conditions; 52 female; age range: 4 months, 1 day – 5 months, 13 days; mean age: 4 months, 20 days). Eight additional infants were excluded for fussiness, inattentiveness, or parental interference.

#### Materials and Procedure

The displays and procedures were similar to those used in Experiments 1a, 2a and 2b, except that the inner members of each group were removed, leaving three characters (Figure 5). The sequence of events was the same. The familiarization events each depicted an interaction between the central character and one of the two side characters. The test events also followed those used in previous experiments, with a slight alteration in the endpoints of the approach trajectories and the positions of the two characters during the synchronous dancing portion of the event, to adjust for the removal of the third group member (Figure 5c and 5d). The order of jumping within each familiarization event (side or central character first) and the direction of the approach events (central character toward side characters or side characters toward central character) were crossed in a 2 × 2 design to create four conditions, varying orthogonally in whether they depicted responder or target approach, executed by one or two actors. All looking times were recoded by a blind, offline coder; correlation of looking times for the two independent coders was high (*r* = 0.99 across 25% of participants). Seventeen participants had a single trial pair excluded due to the ineligibility of one or more trial from the pair.

#### Data analysis

We began by using one-sample t-tests to compare the proportion of congruent looking across the two responder approach conditions and across the two target approach conditions to chance (50%), investigating whether either condition type elicited the previously observed preferential attention to congruent approach trials. Two ANOVAs, one for responder approach conditions and one for target approach conditions, compared the results of Experiment 4 to earlier experiments, assessing the between-subjects effects of number of characters (3 vs. 5), the number of actors at test, familiarization order, and test order on congruent looking preferences in the responder approach and the target approach conditions, respectively.

Two final analyses were conducted, parallel to the central analyses of Experiments 1 and 2. First, one-sample t-tests compared the proportion of looking to congruent trials to chance in each of the four conditions of Experiment 4. Second, an ANOVA including the between-subjects factors of the type of test trial (responder vs. target approach), number of test actors (one vs. two), familiarization order (imitation first or second) and test order (congruent approach first or second), tested whether the pattern of looking to congruent trials varied across the four conditions of Experiment 4.

### Results

#### Responder approach conditions

Infants in the responder approach conditions showed no looking preference for the congruent test trials (49.8%, *t*(47) = 0.07, *P* > 0.9; Figure 6a and 6c), in marked contrast to the pattern observed in the previous five-character experiments. The ANOVA comparing the two responder approach conditions of Experiment 4 with the corresponding conditions of Experiments 1 and 2 revealed a main effect of the reduction from five to three characters (*F*(1,64) = 6.24, *P* < 0.05), as well as an interaction between that factor and the number of actors at test (one vs. two individuals or groups) (*F*(1,64) = 4.77, *P* < 0.05). Infants showed less of a preference for congruent responder approach events in the three-character context of Experiment 4 than in the five-character context of Experiments 1 and 2, but this difference was primarily driven by the condition presenting test events in which a single actor affiliated with two targets (Figure 6a).

**Figure 6.**
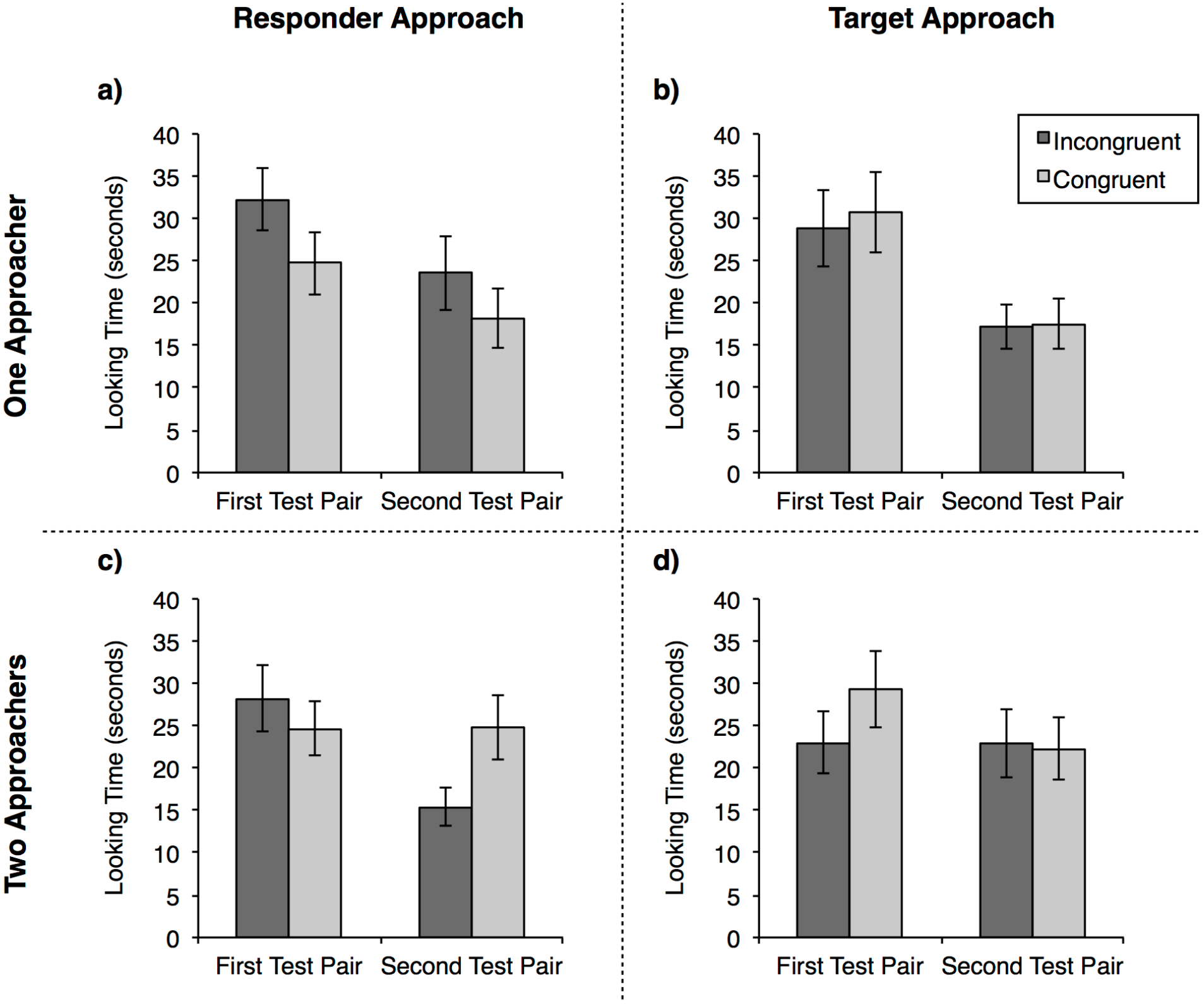
Mean looking times for each test pair from the four conditions of Experiment 4. Responder approach conditions are represented on the left and target approach conditions on the right. Conditions with a single approacher are represented on the top row, and those with two approachers on the bottom row. Error bars represent SEM.

The analyses of individual conditions confirmed these patterns. Infants who observed two responders alternately approach a single target showed weak, non-significant looking preferences between the two events (*M* congruent looking = 54.8%, *t*(23) = 1.33, *P* > 0.1; Figure 6c), whereas infants who observed a single responder alternately approach two targets showed a non-significant trend toward longer looking at the *incongruent* event (*M* congruent looking = 44.9%, *t*(23) = 1.82, *P* = 0.08; Figure 6a). Looking preferences differed significantly across these two conditions: *t*(46) = 2.17, *P* < 0.05). Thus, infants’ attention to the congruent events decreased with the reduction from five to three characters, and this decrease produced a reversal in the direction of looking preferences when the number of actors in the test events decreased as well.

#### Target approach conditions

Infants in the target approach conditions did not differentiate the congruent and incongruent test events of Experiment 4 (*M* congruent looking = 51.2%, *t*(47) = 0.59, *P* > 0.5; Figure 6b and 6d). The ANOVA comparing Experiment 4 to previous target approach conditions from five-character experiments revealed no evidence that reducing the number of characters impacted infants’ relative looking to the congruent versus the incongruent trials (*F*(1,64) = 0.41, *P* > 0.5). There were no effects of or interactions with the number of actors at test (all *P* > 0.5). Moreover, comparisons of the proportion of congruent looking to chance revealed no effects in either the one-actor condition (*M* congruent looking = 51.7%, *t*(23) = 0.56, *P* > 0.5; Figure 6b) or the two-actor condition (*M* congruent looking = 50.7%, *t*(23) = 0.25, *P* > 0.8; Figure 6d), which did not differ from each other. In sum, no matter how the displays were simplified, infants show no evidence of predicting affiliation by actors or groups that had been the targets of imitation.

#### Four-condition analysis

The ANOVA comparing the four conditions of Experiment 4 revealed a strong main effect of test order (*F*(1,80) = 15.13, *P* < 0.001). Across all conditions, a greater proportion of looking time was directed to congruent events when they were presented first (56.1%) as opposed to second (44.9%). Though there was an interaction between condition and test order in Experiment 2b, this was the first experiment in which such a robust order effect was observed. It may reflect greater attentiveness from infants at test, due to the elimination of two characters and their associated actions during the familiarization events, and thus a larger reaction to the novelty of the approach and synchronous motion presented in those events.

With respect to the potential difference in expectations of affiliation by imitators vs. their targets, there was no main effect of responder vs. target approach (*F*(1,80) = 0.21, *P* > 0.6), and no significant interaction between the type of approach trial and the number of actors at test (one vs. two: *F*(1,80) = 3.52, *P* = 0.064). The non-significant trend in this interaction nevertheless accords with the finding, described above, that infants in responder approach conditions displayed different looking preferences when tested with one vs. two actors, while infants in the target approach conditions were not sensitive to this factor. Thus, although the overall patterns of Experiment 4 were inconclusive in themselves, the analysis suggests that patterns of attention observed in the responder approach conditions continued to be sensitive to factors affecting the complexity of the displays, whereas looking times collected from the target approach conditions still failed to show any evidence of differentiation between congruent and incongruent trials or any impact of task complexity.

### Discussion

The findings of Experiment 4 were not significant when considered alone. When compared to the findings of Experiments 1 and 2, however, they help to adjudicate between competing accounts of the preferential attention toward congruent responder approaches observed in those experiments. The presentation of less complex versions of the displays from the earlier experiments substantially reduced the congruency preference observed in the experiments presenting five characters: a reduction that is consistent with stronger or higher confidence expectations regarding imitators’ affiliative behavior. Thus, the findings accord with the hypothesis that infants do base expectations about others’ likely affiliative behavior on observations of their imitative or non-imitative responses toward interaction partners, and that the complexity of the observed interaction affects the strength of those expectations.

Experiment 4 provides suggestive support for a second prediction that follows from the hypothesis that the complexity of observed social interactions influences the strength of infants’ expectations that imitators will affiliate with their targets: infants appear to draw stronger inferences about the affiliative choices of a single responder toward two different targets than about the affiliative choices of two different responders toward the same target. Comparing across the two conditions testing expectations of affiliation by imitators, infants showed a significantly greater preference for the congruent test event in the scenario that presented affiliative actions by two different actors than in the scenario presenting two affiliative actions by a single actor. This finding is consistent with the thesis that infants expect imitators to affiliate with their targets, and that this expectation leads to looking preferences in opposite directions, depending on the difficulty of encoding these social events.

Despite the evidence that the reduction from five to three characters, or from two to one affiliative actors, made the present events easier to process, these decreases in complexity had no effect on infants’ relative looking to congruent and incongruent target approach trials. As in the five-character experiments, infants in Experiment 4 displayed no signs of differentiating events in which characters approached interaction partners who had imitated them versus ones who had not. This consistent lack of differentiation in the target approach conditions suggests that young infants’ failure to look longer at congruent test events in the target-approach conditions of Experiment 2 is not attributable to the cognitive demands that those conditions posed being just outside of participants’ capacity. Instead, it appears that either (1) generating expectations of targets’ affiliative behavior toward responders on the basis of imitation is too complex for young infants even in these highly simplified displays (reducing the likelihood that such an expectation would operate in real world social cognition), or (2) at this age, infants’ understanding of social interaction does not include any relationship between imitative behavior and the attitude of the target of that behavior toward the imitator. This asymmetry between expectations for the agent versus the target of imitative behavior has implications for understanding the nature of infant social cognition and its potential relationship to infants’ responses to and engagement in first person imitation. We consider these implications in the general discussion.

Nevertheless, the results of Experiment 4 raise two concerns. First, infants may not make *any* inferences about affiliation based on imitation observed in dyadic, as opposed to group, contexts. This possibility is consistent with the finding that the proportion of congruent looking did not differ significantly from chance (in either direction) in any of the four conditions of Experiment 4: the only significant effect in the experiment, considered by itself, came from the comparison of looking patterns in the two responder approach conditions (one affiliative actor vs. two). Second, since infants in the responder approach/two actor condition still spent, on average, more time looking to congruent than to incongruent trials, the inference that reducing complexity decreases infants’ relative bias to attend to imitation-congruent approach events is based primarily on the lone condition in which a single responder was shown approaching the targets it did and did not imitate. Given the importance of the non-significant incongruency preference observed in this condition, it bears further investigation. In Experiment 5, we attempt to replicate and strengthen the effect in the single condition within this series in which 4-month-old infants exhibit any trend toward an incongruity preference.

## Experiment 5

In Experiment 5, we ask whether 4- to 5-month-old infants, presented with imitative and non-imitative interactions for a longer period of familiarization prior to test than those in the preceding experiments, look longer at events in which a character approaches a lone target that it did *not* imitate: the incongruent test event. To this end, infants were shown exactly the same events used in the responder approach/one actor condition of Experiment 4, with one change in method. In an effort to enhance infants’ memory for the imitative interactions, and to reduce the strong order effect observed across all conditions of Experiment 4, we altered the order of the events such that all eight familiarization events preceded the four test events, which were now presented at the end of the experiment.^6^

### Methods

#### Participants

Participants were 24 4- to 5.5-month-old infants (8 female; age range: 4 months, 5 days – 5 months, 16 days; mean age: 4 months, 28 days). Three additional infants were excluded for fussiness or inattentiveness.

#### Materials and Procedure

The materials and procedure were identical to those used in the responder approach/one actor condition of Experiment 4, except that the sequence of events was altered so that infants first saw all eight familiarization events and then all four test events. The familiarization events alternated between imitative and non-imitative interactions and the test events alternated between congruent and incongruent events, with the order of these alternations (imitation first or second; congruent first or second) counterbalanced across participants. There were four infant-directed pauses during the familiarization phase, one after every second event, lasting up to 60 s or until infants looked away for 2 consecutive seconds, as determined by a blind, online coder. All looking times were recoded by one or two blind, offline coders; the correlation between looking times recorded by these two coders was high (*r* = 0.96 across 25% of participants). One pair of trials each was excluded for three different participants.

#### Data analysis

We began by comparing the total looking time (in seconds) to all valid test trials in this experiment to that in the comparable condition of Experiment 4 with an independent samples t test, to assess the effect of positioning all test trials at the end of the display on infants’ overall attention. Then, as in previous experiments, the proportion of looking to congruent trials, calculated first within each test pair and then averaged across the two pairs, was compared to chance (50%) by a one-sample, one-tailed t-test, with the directional prediction that infants would look longer at the *incongruent* event.

Then a two-way ANOVA tested for effects of imitation order or test order on proportion of looking to congruent trials. An independent-samples t test compared the findings of Experiment 5 to those of the corresponding condition of Experiment 4, to assess whether infants’ looking preference for the incongruent event increased with the more concentrated period of familiarization. Finally, a one-way t-test compared the proportion of looking to congruent trials across the two conditions to chance, to get an estimate of the strength of the effect of congruency on infants’ relative looking to the present test events.

### Results

The change in event sequence led participants in this experiment to spend less total time looking at all test trials (54.05 s) compared to participants in corresponding condition of Experiment 4 (95.72 s; *t*(46) = 3.12, *P* < 0.01). Despite this reduction in overall looking, infants directed a larger proportion of looking time to the incongruent test trials, replicating the effect that appeared as a nonsignificant trend in Experiment 4 (*M* looking to congruent events = 46.3%, *t*(23) = 2.03, *P* < 0.05; Figure 7). The two-way ANOVA testing for effects of the between-subjects factors of imitation order and test order on proportion of looking to congruent trials revealed no main effects or interactions (all *P* > 0.05). The analysis comparing the test trial looking preference in Experiment 5 with that of the corresponding condition of Experiment 4 revealed no significant increase in the strength of the incongruity preference with the presentation of an uninterrupted string of familiarization trials (*t*(46) = 0.42, *P* > 0.6). Across these two experimental conditions, infants showed a moderately strong looking preference for the incongruent event in which the responder approached the target it did not imitate (54.4%, *t*(47) = 2.66, *P* = 0.01, Cohen’s *d* = 0.38).

**Figure 7.**
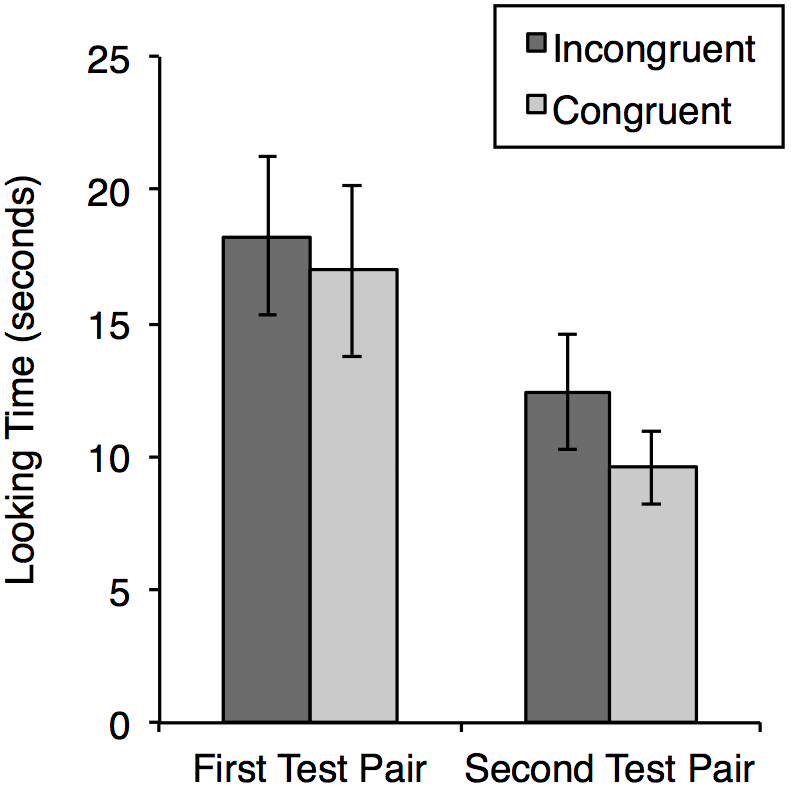
Mean looking times for each test pair in Experiment 5, featuring a single responder and responder approach test trials. Error bars represent SEM.

### Discussion

In Experiment 5, infants looked longer when an individual made an inconsistent approach toward the social partner it did not previously imitate than when that same individual approached the imitated partner. This finding provides evidence that infants base inferences about others’ social affiliation on observations of their imitative and non-imitative responses to different social partners. Nevertheless, although this looking pattern attained conventional levels of significance only in Experiment 5, it was no stronger in Experiment 5 than in the corresponding condition of Experiment 4, and the effect size for the incongruency preference across the two experiments is appreciably smaller than the effect size observed in Experiments 1a and 1b, where five-character versions of the same events elicited a strong preference in the opposite direction. The task of generating expectations about third-party affiliative behavior from prior observation of patterns of social imitation may stand at the limit of the abilities of four-month-old infants.

## General Discussion

Five experiments probed the early development of an understanding of social imitation in third party contexts. Guided by the hypothesis that imitation is a highly reliable signal of social attention, motivation, and commitment, we asked whether infants are sensitive to this signal when they view imitative interactions as third parties, whether they expect imitators to choose to affiliate with their former targets rather than with former non-targets, and whether their expectations of imitators are distinct from their expectations of targets. To address these questions, we conducted five experiments, involving over two hundred infants who were tested in eleven different conditions that varied (1) the modality that distinguished imitative and non-imitative interactions (sound vs. movement), (2) the imitative roles of the affiliative actors in the test events (imitators vs. targets of imitation), (3) the composition of the social parties (individuals vs. pairs, resulting in a total of 3 or 5 characters), (4) the number of different affiliative parties engaging in approach at test (one vs. two), (5) the number of different actions that distinguished some characters from others (one vs. two), and (6) the age of the participants (4 vs. 12 months). These conditions sought to test not only *if* observations of imitation would affect infants’ attention to subsequent affiliative events but also via what mechanism they might do so.

The principal findings of each condition of these experiments are recapitulated in Figure 8. They are complex, because several conditions revealed systematic preferences that are opposite in direction to those that one would predict under the assumptions that infants look longer at unexpected events, and that infants view imitation as a reflection of affiliation. In the literature on perceptual and cognitive development, however, such preference reversals typically occur when the ability that one is testing for is present but fragile, because the test places high cognitive demands on the infants and lowers their confidence in their interpretation of the events presented to them. Numerous aspects of the present findings are consistent with the observed cases of congruency preference reflecting this phenomenon, including the change with age from preference for the congruent to the incongruent events, and the reduction in the congruity preference with manipulations that reduced task demands.

**Figure 8.**
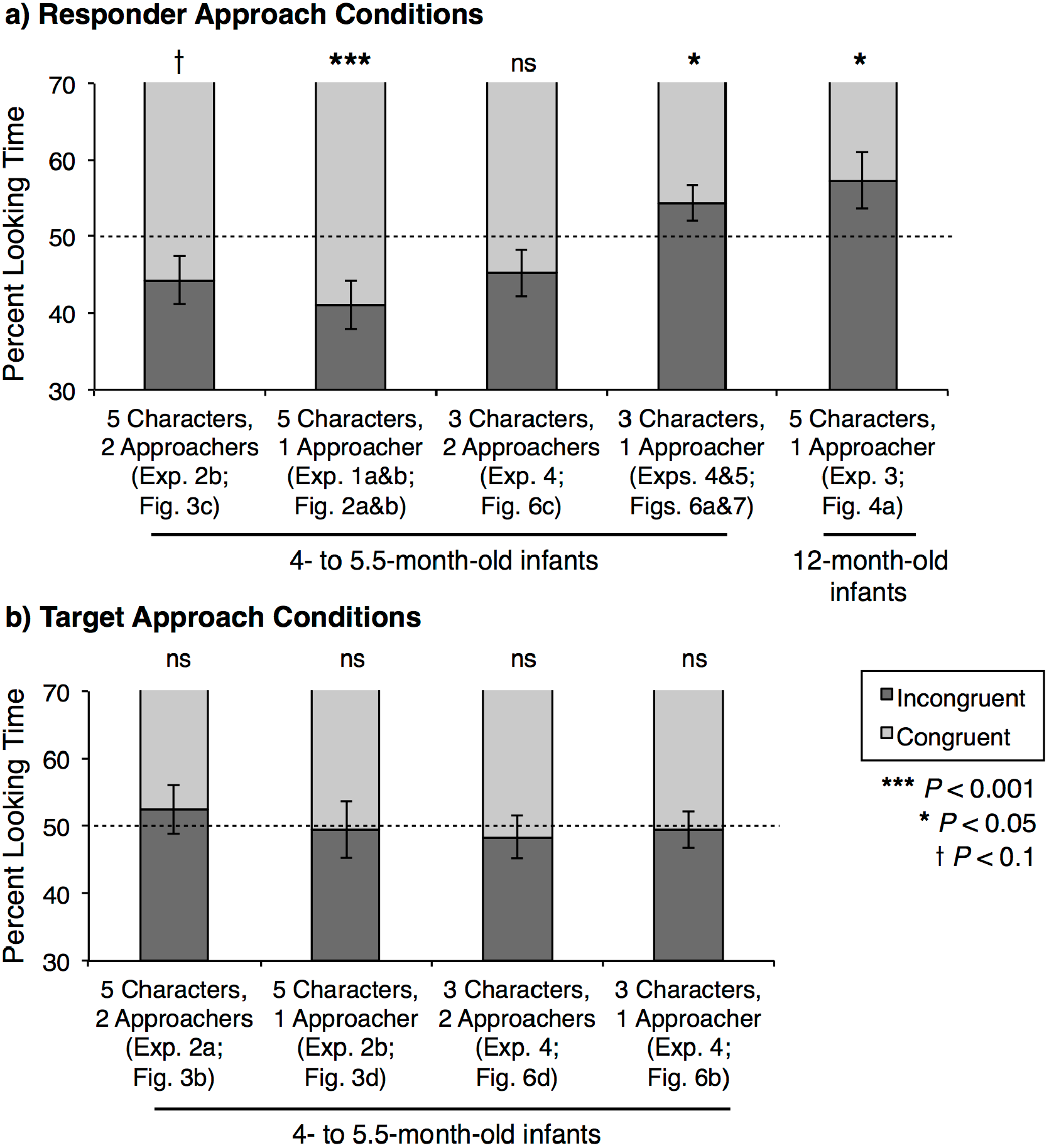
Proportional looking time to (a) Responder Approach and (b) Target Approach conditions. Responder Approach conditions show a shift from congruency preferences to incongruency preferences with decreasing complexity and increasing age, while Target Approach conditions show no sensitivity to complexity. For bars representing two experiments, significance values are drawn from one-sample t-tests that pooled data from both experiments and compared percent looking to congruent trials to chance. Error bars represent SEM.

Together, therefore, the present findings support three conclusions. First, infants as young as 4 months expect a character who has imitated one of two other characters to approach the target of his imitation, and they show no comparable expectations about a character who has been a target of imitation. Second, this expectation is fragile, as looking preferences reverse in direction with increases in the number of actors in the imitative interactions, the number of actors at test, or the age of the infants. Third, young infants’ response to imitation cannot be explained solely by learning about the low-level features of common imitative acts, because it was exhibited in response to abstract animated characters whose sounds and motions were unlike any vocalizations and biomechanical motions that humans perform. These three findings seem uniquely consistent with the thesis that young infants view imitation as reflecting the social attention and orientation of the imitator, but do not speak to the attention or motives of the target of imitation. Acts of imitation may be among the first social signals that young infants understand.

The contrast between infants’ performance in the experimental conditions presenting approach by responders and approach by targets is particularly striking, because the experimental conditions testing infants’ responses to approach by imitators and by targets of imitation presented exactly the same events during familiarization and during test; only the pairing of these events differed. Moreover, these pairings yielded responder approach and target approach conditions that differed only in the ordering of actions during familiarization. Although changes in ordering sometimes affect the detectability or memorability of events, such effects cannot plausibly account for the present findings. If infants detected and retained representations of the actions of imitators but not targets because the imitative actions benefitted from a recency effect, for example, then infants would have had no basis for assessing the similarity between the responders’ and targets’ actions, and thus no basis for differentiating congruent and incongruent approach trials in any condition. Moreover, if infants had a strong bias to remember the most recent information at test, we would have expected to see effects of familiarization order, with infants perhaps showing stronger expectations for imitators to approach their targets (or vice versa) when imitative interactions, rather than non-imitative interactions, directly preceded the test events. No such order effect was observed. The asymmetry between the data from the responder and target approach conditions instead seems to reflect a robust asymmetry in the infants’ inferences about the affiliative dispositions of responding social partners versus those of their targets.

These findings suggest that imitation has social significance for infants, at an age when infants’ own imitative skills are limited. They also provide new insight into infants’ early social cognitive capacities. The behavior exhibited by infants requires three basic aspects of social cognition. First, infants must have noticed when interaction partners’ behaviors were similar rather than different. Second, they must have tracked each party’s role in the interaction, encoding which party initiated the interaction and which party responded. Finally, infants must have attributed an attitude or behavioral disposition on behalf of one party toward the other.

Prior to this research, it was not clear that infants of this age were capable of any of these cognitive achievements. Although young infants are frequently imitated by their parents, there is little evidence that they recognize imitative responses, over and beyond merely contingent ones, until the second half of the first year. The present third-party design required only that infants match two observed actions, not that they match an observed action to an executed one; this may have aided infants’ detection of imitation. With respect to tracking separate parties’ roles in social interactions, some research provides evidence that young infants differentially evaluate actors who help versus hinder a third individual (Hamlin, Wynn & Bloom, 2010). It is not clear, however, whether infants under 6 months responded to the helper or hinderer’s distinctive role in that interaction or only to their participation in it.

Finally, researchers have repeatedly failed to find evidence that young infants form expectations about individuals’ social attitudes or behavioral dispositions under circumstances where infants closer to one year of age succeed. For instance, infants under 10 months do not expect others to approach those who have helped rather than hindered them (Kuhlmeier et al., 2003; Hamlin et al., 2007), nor do they predict socially dominant or subordinate behaviors based on size, as older infants do (Thomsen et al., 2011). There is evidence that infants between 8 and 12 months of age base expectations of shared behavior on social affiliation and vice versa (Powell & Spelke, 2013; Liberman et al., 2016), but this does not require the attribution of one individual’s social disposition toward another. In contrast, the current experiments demonstrate that even infants under 6 months of age consider social actions to be evidence of the actor(s)’ attitude or behavioral disposition toward the target(s) of the actions, and use that inferred disposition to guide expectations about the actor’s novel social behaviors toward the same target.

The current results therefore extend and shed light on previous research on young infants’ social cognition. The null results in our target approach conditions are consistent with young infants’ failure to expect the recipients of helpful acts to preferentially approach helpers. The success of our responder approach conditions raises the possibility that young infants would expect helpers to be more likely than hinderers to approach the target of their helpful actions, and that they may expect an individual to approach someone they have helped rather than someone they have neglected or hindered, but to our knowledge no experiments have tested these hypotheses.

We also are unaware of any published evidence that infants as young as 4 months of age prefer those who imitate them, as do older infants, children, and adults (Meltzoff, 1990; Agnetta & Rochat, 2004; Thelen et al., 1975; Chartrand & Bargh, 1999). Even if a preference for imitators does extend to the first few months of life, however, this does not automatically imply that infants will attribute similar preferences to the targets of imitation that they observe as third parties. There are a number of factors that may prevent young infants from extending this attribution in the present experiments. For example, because the target characters in these experiments show no social responses to the actions of other characters during the imitative interactions, infants may lack information as to whether the target character itself is a social being with social dispositions. The literature on infants’ understanding of helping and hindering is again instructive, as it has found that infants under 10 months show preferences for helpers over hinderers but do not expect the target of the helping and hindering acts to show a similar preference (Kuhlmeier et al., 2004; Hamlin et al., 2007).

Regardless of the source of young infants’ failure to expect the targets of imitative (or helpful) social acts to approach those actors, older infants may come to expect targets to approach those who imitated them over those who did not, consistent with their expectation of targets’ approach toward helpers over hinderers and with older children and adults’ use of imitation to elicit liking from others (Over & Carpenter, 2009; Watson-Jones et al., 2016; Lakin et al., 2008). Indeed, such a development could relate to young infants’ own first person engagement in imitation, which is severely limited in the first six months of life (Ray & Heyes, 2010; Anisfeld, 2005). If infants increasingly grasp the positive effect of imitation on the target’s reciprocal attitude toward the imitator as they approach their first birthday, this could partly explain infants’ increasing engagement in voluntary social imitation around that age (e.g. Jones, 2007; Carpenter, Nagell & Tomasello, 1998).

The current data suggest that the inferences infants drew from imitators’ repetition of their targets’ behavior was social in nature. Although general attention to repetitive actions or agents could explain greater attention on infants’ own part toward imitating rather than non-imitating responders, it is unclear what domain-general explanation could account for the selective expectation that a repetitive entity would move toward the original exemplar of its behavior but not vice versa. In contrast, this selective expectation can be expressed easily as the tenet of a naïve social theory: social actions are evidence of the actor(s)’ attitude toward the target(s) of the actions. In the case of social imitation specifically, the present findings provide evidence that infants infer that imitators have a prosocial stance toward those whom they imitate that can guide expectations of behavior in further social interactions. These findings thus suggest continuity in the perception of imitation as a positive social behavior, a perception shared by older infants, children and adults (Meltzoff, 1990; Agnetta & Rochat, 2004; Carpenter et al., 2013; Over & Carpenter, 2015; Chartrand & Bargh, 1999).

The present findings raise many questions for future research. In particular, our experiments do not reveal whether infants view imitation as reflecting only social attention, or also stronger and more specific behavioral dispositions, such as a disposition to help or cooperate with the target of their imitation. They also do not reveal whether infants attribute mental states to those who imitate others, such as a liking for the target of their imitation (Over & Carpenter, 2015) or a belief that the target is more powerful, presitigious or skilled (Henrich & Gil-White, 2001; Chudek, Heller, Birch & Henrich, 2012)?

Our experiments also do not clarify what range of actions infants interpret as social signals when they view imitative interactions. We tested infants’ responses to social events involving imitation of sounds or motions that were emitted in contexts involving no objects: actions that occur during social communication such as speech, emotional vocalizations, and gestures. Would infants attribute social motives to imitators of a broader class of actions, including instrumental, object-directed actions or involuntary actions? Answers to this question may help to reveal whether infants view imitative behavior specifically as a type of communication or more generally as a type of social behavior.

Beyond the study of imitation, the present methods and findings may serve to address more general questions regarding the early development of social knowledge. They provide methods that could serve, in future investigations, to determine whether infants consider social actions to be evidence for stable social preferences on the part of the actor, or as momentary communicative behaviors, through experiments that test for the consistency of infants’ expectations of affiliative behavior by imitators over time. Their methods also could serve to investigate whether infants view social actions as informative about the larger social landscape, through experiments testing whether infants make transitive inferences from a pattern of imitative acts, expecting imitators to copy and preferentially engage with their targets’ social affiliates. Finally, their methods could investigate the development of infants’ understanding of dominance, competence, or prestige, through experiments testing whether infants expect a new social character to imitate a target who previously had been imitated by others.

However these questions are answered, the present findings have implications for our broader understanding of the early social cognitive capacities of very young infants. Substantial evidence demonstrates that young infants evaluate others on the basis of their behavior: infants prefer some social partners over others based on the language they speak, their tone of voice, and their social actions (Kinzler et al., 2007; Schachner & Hannon, 2011; Hamlin et al., 2007). The present experiments provide the first evidence, however, that infants below six months can go beyond their own personal evaluations and make inferences about the attitudes that others have toward their social partners. Moreover, the asymmetry we observe between infants’ expectations about imitators and about targets of imitation shows that infants can go beyond inferences of mutual affiliation or shared group membership (e.g. Liberman et al., 2014; Powell & Spelke, 2013) and attribute selective social attitudes to the individual participants in a social interaction. Early in development, therefore, human minds are equipped with a system of social inference. This system likely supports very young infants’ learning about the social partners and social actions that surround them, fostering their social cognitive development. If that is the case, then experiments focused on young infants’ third party social inferences should provide a fruitful approach both for further elucidating young infants’ understanding of their social world, and for probing the sources of our distinctively human social minds.

## Acknowledgements

We thank Ellyn Schmidt and Natasha Kalra for assistance with data collection and coding. This research was supported by a grant to ESS from NIH (HD23103), by a grant from the Simons Foundation to the Simons Center for the Social Brain, and by the Center for Brains, Minds and Machines (CBMM), funded by NSF STC award CCF-1231216.

## Supplementary Information

### Log transformed looking time analyses

Prior to data collection, we planned to analyze proportional looking time, as it equally weights differences in looking to contrasting trial types across pairs of test trials that frequently differ in overall looking time as infants’ attention to displays decreases over the course of an experimental session. Subsequent to data collection, Csibra et al., (2016) published reanalyses of a large number of looking time experiments, recommending analyses of log-transformed looking times for violation of expectation studies such as the ones presented here. They argued that the log transform helps to correct skewed distributions common in looking time data but not necessarily detectable in samples of the size recruited for a typical experiment (Csibra et al., 2016). Although our original proportion scores may also have helped to reduce skewness, both by placing a boundary on the extent of the right tail of the distribution and by moving cases of consistently short looking times across both trial types to the middle of the distribution, we decided to also reanalyze our data in accord with these recommendations. These analyses are presented below. Because the alternating familiarization-test-familiarization-test design of the majority of our experiments results in significantly shorter looking times to trials in the second test pair compared to the first, we continued to exclude both trials from test pairs in which one trial was invalid. As described below, the outcomes of these analyses accorded with those of the primary analyses in nearly all cases.

### Experiment 1a

A paired samples t test revealed significantly longer looking to congruent approaches (1.35) than incongruent ones (1.16; *t*(15) = 2.94, *P* = 0.01). A repeated measures ANOVA with trial type as a within subjects factor and familiarization and test order as between subjects factors found only a main effect of trial type, *F*(1,12) = 7.19, *P* < 0.05. All other main effects and interactions were not significant (*P*s < 0.4). All these findings accord with the primary analyses.

### Experiment 1b

As in Experiment 1a, infants looked significantly longer to congruent approaches (1.36) than incongruent ones (1.19; *t*(15) = 2.50, *P* = 0.024), and a repeated measures ANOVA testing for effects of familiarization and test order revealed only a main effect of trial type (*F*(1,12) = 6.98, *P* < 0.02; all other *P*s > 0.05). A repeated measures ANOVA comparing looking times in Experiment 1b (movement imitation) to those in Experiment 1a (sound imitation) with trial type as a within subjects factor and imitation type as a between subjects factor revealed a main effect of trial type (*F*(1,30) = 14.66, *P* < 0.005), and no evidence of an interaction with the type of imitation displayed (*P* > 0.9). These findings accord as well with the primary analyses.

### Experiment 2a

In accord with the primary analyses, the paired samples t test provided no evidence that infants in Experiment 2a, who observed imitated and non-imitated groups alternately approach the lone responding character, differentiated between congruent (1.30) and incongruent approach trials (1.35; *t*(15) = 0.65, *P* > 0.5), and the repeated measures ANOVA testing for familiarization and order effects revealed no significant main effects or interactions (all *P*s > 0.1). A repeated measures ANOVA comparing looking times to the target approach events of Experiment 2a with those to responder approach events in Experiments 1a and 1b revealed a significant interaction between trial type and approach type, *F*(1,46) = 7.47, *P* < 0.01, in accord with the main effect in the primary analysis.

### Experiment 2b

Separate paired samples t tests for the target approach (congruent looking: 1.25, incongruent looking: 1.25, *t*(15) = 0.04, *P* > 0.9) and responder approach conditions (congruent looking: 1.31, incongruent looking: 1.21, *t*(15) = 1.78, *P* = 0.095) both failed to provide evidence of a difference looking time, though the responder approach condition showed a non-significant trend toward longer looking to congruent trials. Analogous to the ANOVA testing for order effects in the primary analyses, the repeated measures ANOVA testing for effects of trial type (within subjects), approach condition, familiarization order, and test order (all between subjects) found a trial type x condition x test order interaction, *F*(1,24) = 4.90, p < 0.05, reflecting the fact that infants in the responder approach condition looked longer to the congruent events than incongruent events when congruent events were presented first (1.27 vs. 1.00, respectively; *t*(7) = 4.13, *P* < 0.01), but looked for an equivalent amount of time at the event types when incongruent trials were presented first (congruent: 1.31, incongruent: 1.41, *P* > 0.4).

Experiments 1a, 2a, and 2b comprised a 2 x 2 design, varying the roles the paired and lone characters played as responder or target in the interactions, as well as whether the paired or lone characters performed the approach actions at test. The repeated measures ANOVA testing for effects of trial type (within subjects) as well as responder identity, approacher identity, and test order (all between subjects) across these four conditions revealed a significant trial type x responder identity x approacher identity interaction, *F*(1,56) = 5.81, *P* < 0.05, reflecting the fact that infants looked longer to the congruent approach events than incongruent ones when responder and approacher identity matched, (congruent: 1.33, incongruent: 1.18, *t*(31) = 3.36, *P* < 0.005) but not when the target(s) were the one(s) approaching (congruent: 1.28, incongruent: 1.30, *t*(31) = 0.66, *P* > 0.4). All these findings accord with those of the primary analyses.

### Experiment 3

A paired samples t test found evidence in support of the one-tailed prediction that, in contrast to 4-month-old infants, 12-month-old infants would look longer at incongruent approaches (1.19) by a responder toward the targets it did not imitate, relative to congruent approaches toward imitated targets (1.04; *t*(15) = 2.05, *P* = 0.029). A repeated measures ANOVA including test order as a between subjects factor found a main effect of test order, *F*(1,14) = 6.04, *P* < 0.05, reflecting longer overall looking for infants who saw incongruent approaches first, but no interaction between test order and trial type.

A repeated measures ANOVA comparing looking time to congruent and incongruent approaches by a lone responder across this experiment and Experiments 1a and 1b, with age group as a between subjects factor, found a significant trial type x age group interaction, *F*(1,46) = 15.79, *P* < 0.001. All these analyses accord with the primary analyses.

### Experiment 4

Experiment 4 comprised a 2 x 2 cross of the role that the approacher(s) had played in the interactions (responder or target approach) and the number of different approaching actors across the test trial types (one or two). Paired samples t tests comparing log transformed looking times to congruent and incongruent trials within each of these four conditions revealed no significant differences, though there was a non-significant trend toward longer looking times to incongruent trials (1.32) than congruent trials (1.21) in the responder approach, one approacher condition (*t*(23) = 1.96, *P* = 0.06; all other *P*s > 0.1). A repeated measures ANOVA with trial type as a within subjects factor and approach type (responder or target), number of approachers (1 or 2), familiarization order and test order as between subjects factors revealed an overall interaction between trial type and test order, *F*(1,80) = 14.44, *P* < 0.001, reflecting longer looking times to whichever trial type came first, in accord with the primary analyses.

A repeated measures ANOVA comparing the responder approach conditions in the current experiment to those in Experiments 1a and 2b, including trial type as a within subjects factor and number of approaching individuals or groups (1 vs 2), overall number of characters (5 vs. 3), familiarization order, and test order as between subjects factors, revealed an interaction between trial type and overall number of characters, *F*(1,64) = 5.924, *P* < 0.05, as in the primary analysis, reflecting a weaker congruency preference when displays were simplified by the use of fewer characters. There was also a three-way trial type x overall number x number of approachers interaction *F*(1,64) = 5.01, *P* < 0.05, reflecting the fact that the reduction in the congruency preference was strongest in the simplest condition, when infants were only asked to attend to a single approacher, moving toward one character it had imitated or one it had not. (Indeed there was a trend toward a preference for incongruent trials in this condition, as noted above.) There was also an interaction between trial type and test order, *F*(1,64) = 11.98, *P* < 0.005, reflecting a stronger relative preference for congruent trials when the congruent trials came first rather than second in the test pairs. There were no between-subjects effects of these factors on overall looking time to the different conditions. All these findings accord with the primary analyses.

An analogous repeated measures ANOVA for target approach conditions from Experiments 2a, 2b, and 4, comparing looking times to congruent and incongruent trials under conditions of different numbers of approachers, numbers of overall characters, familiarization orders and test orders failed to find any evidence of an effect of reducing the number of characters, or of an interaction between this variable and the number of approachers (both *P* > 0.4). This negative finding accords with the negative finding in the primary analysis.

### Experiment 5

A one-tailed paired samples t test showed that looking times to incongruent trials (1.07) were significantly longer than those to congruent trials (1.00; *t*(23) = 2.03, *P* = 0.027), as in the primary analysis. A repeated measures ANOVA with familiarization and test order as between subjects factors revealed a main effect of trial type, *F*(1,20) = 4.44, *P* < 0.05, as well as a three-way interaction between trial type, familiarization order, and test order, *F*(1,20) = 4.38, *P* < 0.05. (There was a non-significant trend [*P* <0.1] toward a similar interaction between familiarization and test order in the proportional analyses that we did not report in the main text.) This interaction reflected a pattern in which infants devoted relatively more attention to the first test trial when it represented a change in order from the preceding familiarization events (e.g. when the imitative interaction preceded the non-imitative one during familiarization, but the approach toward the non-imitated target occurred first, or vice versa). Thus extending the familiarization sequence eliminated the overall order effect observed in Experiment 4, in which infants attended more to the first trial presented in a test pair, but may also have increased infants’ sensitivity to the stable pattern of alternation present across all familiarization trials, leading them to notice when the pattern of interaction order was violated at test.

An additional repeated measures ANOVA comparing looking times in this experiment, featuring blocked familiarization trials, to those in the analogous condition of Experiment 4, which used alternating familiarization and test trials, revealed a main effect of trial type F(1,46) = 7.60, *P* < 0.01 and a main effect of familiarization type on overall looking time to test trials, *F*(1,46) = 10.99, *P* < 0.005, but no interaction between the two factors (*P* > 0.6).

## Footnotes

1 Although the displays were relatively simple, they presented characters with a number of attributes that have been found, in other experiments, to carry social meaning: not only sound and motion, but also shape, color, facial features, emotional expression, and spatial location. To ensure that only one behavior (sound in Experiment 1a and motion in Experiment 1b) appeared to involve imitation of one character, the other attributes either were held constant across all five characters and therefore did not serve to distinguish some characters from others (this was the case for motion in Experiment 1a, and for shape, facial features and emotional expression in both versions of the experiment), or they varied across the five characters and therefore did not serve to connect the responding character to one of the other two characters more than to the other character (this was the case for color and spatial location in both versions of the experiment).

2 Because the proportion measure fails to capture differences in absolute levels of looking time, we also report mean looking times to the congruent and incongruent events (see Figure 2), and we analyzed the data by ANOVAs based on these looking times, log-transformed to correct for positive skew (after Csibra, Hernik, Mascaro, Tatone & Lengyel, 2016). The findings of these analyses are reported in the Supplementary Information, and are in broad accord with the findings of the principal analyses.

3 Experiment 3 was conducted at about the same time as Experiments 1 and 2, but was conceived as an independent investigation undertaken to explore the conditions under which infants would expect an individual to be accepted versus excluded by a social group. There are some minor differences in displays, described below, resulting from this historical circumstance, as well as the tailoring of the displays to two different age ranges. However, none of these differences alter the overall logic of the displays, in which a lone character imitates the actions of one group and not another, and then alternately approaches and acts in synchrony with each of the groups.

4 It is possible that the 12-month-old infants found it easier to perceive and remember the imitative interactions not because they are older, but because both the motion and sound produced by the imitative and non-imitative actors respectively matched and did not match the initiator. If so, then this feature of the displays in Experiment 3, rather than the change in participant age, would have produced all or part of the reversal in infants looking preferences relative to Experiment 1. This possibility is also consistent with the graded expectations hypothesis, as that hypothesis predicts that looking preferences for incongruent test displays will increase with increases both in the maturity of the participants and in the simplicity of the displays: both factors should affect the degree of encoding and strength of subsequent expectations, and have been found to do so in past research using similar characters and imitative events (Powell & Spelke, 2013). However we caution against the assumption that imitative interactions that simultaneously vary both motion and sound will be easier to differentiate for young infants. Tests of the intersensory redundancy hypothesis have found that when there is overlapping information in multiple modalities, young infants’ attend to amodal properties of a stimulus, such as the rhythm or repetition of synchronous auditory and visual input, rather than to specific unimodal features such as those that characterized the imitative interactions in our study (Bahrick, Lickliter & Flom, 2004). Thus mapping differences in multiple unimodal properties to different actors when the amodal properties of rhythm and repetition continue to be similar across all actors may be as demanding as mapping differences that occur in a single modality. Indeed, variation only in one modality, as in Experiments 1 and 2, may help to draw young infants’ attention back from the processing of amodal features to the processing of the motions or sounds that characterize unimodal imitative interactions.

5 The five-character experiments varied similarly, but were confounded by the fact that “one-actor” conditions presented the actions of an individual, while “two-actor” conditions presented the actions of groups. Moreover, the data were inconclusive. Relative attention to congruent and incongruent trials did not vary significantly in one- vs. two-actor conditions, but target approach conditions may already have been too complex for infants. In the responder approach conditions, the difference in the size of the congruency preference in test events presenting one actor (Experiments 1a and 1b) versus two actors (Experiment 2b) may have failed to reach significance due to a lack of power for detecting small effect sizes, a concern addressed by the 50% increase in sample size in Experiment 3.

6 We considered but rejected the idea of running Experiment 5 as a full, infant-controlled habituation experiment, because such experiments are not practical when infants must view an extended series of events within each trial in order for that trial to be valid.

